# Autophagy repression by antigen and cytokines shapes mitochondrial, migration and effector machinery in CD8 T cells

**DOI:** 10.1101/2024.06.10.598276

**Authors:** Linda V. Sinclair, Tom Youdale, Laura Spinelli, Milica Gakovic, Alistair J. Langlands, Shalini Pathak, Andrew J.M. Howden, Ian G. Ganley, Doreen A. Cantrell

## Abstract

Autophagy is important for CD8 T-cells but autophagy timing, triggers and targets are poorly defined. Herein, we show naïve CD8-T cells have high autophagic flux and identify an autophagy checkpoint whereby antigen receptor engagement represses autophagy by regulating amino acid transporter expression and intracellular amino acid delivery. Effector cytotoxic T cells with high levels of amino acid transporters driven by proinflammatory cytokines have low autophagic flux but rapidly reinduce autophagy when amino acid restricted. A census of proteins degraded and fuelled by autophagy shows how autophagy shapes CD8-T cell proteomes. In cytotoxic T-cells, dominant autophagy substrates include cytolytic effector molecules, amino acid and glucose transporters. In naïve T-cells mitophagy dominates and selective mitochondrial pruning supports the expression of molecules that coordinate T-cell migration and survival. Autophagy thus differentially prunes naive and effector T-cell proteomes and is dynamically repressed by antigen receptors and inflammatory cytokines to shape T-cell differentiation.

## Introduction

CD8 T lymphocytes are essential effector cells for adaptive immune responses against viruses and tumours. The proliferation and differentiation of CD8 T cells is directed by their transcriptional programs that are shaped by regulated changes in protein synthesis and degradation. In naïve and memory T cells there are low but critical levels of protein synthesis that sustain expression of transcription factors, transcriptional machinery, metabolic machinery and essential cytokine receptors and adhesion molecules that position the cells for rapid responses to pathogens. The immune activation of T cells is then accompanied by large increases in mRNA translation and protein synthesis rates that implement T cell transcriptional programs to drive T cell clonal expansion and the production of key immune effector molecules.

Protein synthesis is critically dependent on the availability of amino acids. These can be obtained from the environment via membrane amino acid transporters or via macro-pinocytosis of environmental proteins that are subsequently degraded. Amino acids can also be supplied via autophagy, a recycling process that degrades intracellular proteins to provide amino acids for metabolic processes. Autophagy is critical for CD8 T cells since deletion of autophagy regulatory proteins Atg3, Atg5, Atg7 or VPS34 reduces naïve CD8+ T cell survival ^1–7^ and prevents CD8 memory T cell formation ^7–10^. Autophagy also restrains CTL effector function and autophagy loss in effector T cells increases CD8 T mediated anti-tumour immunity ^11,12^.

The importance of autophagy for CD8 T cells makes it essential to understand what signals regulate autophagy and what proteins are degraded and fuelled by autophagy in different CD8 T cell populations. In this context, autophagy is important for quality control of cellular proteomes but protein degradation by autophagy is energetically expensive; degradation must be balanced by protein renewal to ensure cell homeostasis. Initial studies proposed that immune activation of naïve T cells switches on autophagy ^1,2,13–15^. These conclusions are based on assays that measure the presence rather than the activity of autophagosomes. For example, microscopy or western blotting to measure the lipidation of microtubule-associated protein light chain 3 beta-I (MAP1LC3b, LC3b), a classic marker of autophagosomes ^16–18^ or measuring LC3b accumulation following treatment of cells with autophagy inhibitors ^15^. In contrast, experiments using a dynamic autophagy flux reporter model ^7^ revealed that autophagy flux is downregulated in proliferating T cells in vivo and increases as T cell effectors transition to memory cells ^7^.

To re-examine autophagy regulation during T cell activation, the present work uses quantitative proteomics to explore comprehensively how expression of autophagy machinery changes during T cell differentiation. We also use a dynamic autophagy flux reporter to identify the timing, triggers, and regulators of autophy in CD8 T cells responding to immune activation. Additionally, we use mass spectrometry to census T cell proteomes to determine how autophagy shapes naïve and effector CD8 T cell phenotype and functional capacity. This work revises concepts about autophagy control in T cells and shows that autophagy flux is high in quiescent naïve T cells but rapidly repressed by antigen receptor engagement. Immune activated T cells accumulate autophagy machinery, but autophagy flux in effector cells is supressed by proinflammatory cytokines although not by cytokines that drive T cell memory. The ability of T cells to control autophagy flux is shown to be determined by amino acid supply. The control of amino acid transporter expression by antigen receptors and cytokines is thus identified as a critical autophagy checkpoint for T cells because this is what determines the intracellular delivery of amino acids. The current data also map autophagy substrates and proteins fuelled by autophagy in naïve and effector CD8 T cells which informs why precise regulation of autophagy is important for CD8 T cells to initiate and curtail effector function.

## Results

### Immune activated T cells accumulate autophagy machinery but repress autophagy flux

The lipidation or stabilization of the ATG8 protein LC3b (Map1LC3b) has been used to monitor autophagy in many lymphocyte studies ^1,6,7,10,15,19,20^. To assess whether immune activation changes the abundance of LC3b we interrogated high resolution mass spectrometry data that quantitatively analysed T cell proteomes (Howden et al. 2019; Brenes et al. 2023). These data show that MAP1LC3b is not detected in naïve T cells but is abundant as CD8 T cells respond to cognate antigen and differentiate to effector cytotoxic cells (Fig 1A). MAP1LC3b is one of 6 mammalian ATG8 family members and proteomic data revealed that CD8 T cells also increase the abundance of the ATG8 family proteins GABARAP and GABARAPL2 as they differentiate to CTL (Supp Fig 1a). The abundance of other core components of the autophagy machinery also increased as T cells differentiate to CTL including ATG5 and ATG7 and WIPI2 (Fig 1B, 1C, S1A and Supp Table 1). Sequestosome 1 (p62/SQSTM1): a selective autophagy adapter that targets proteins for autophagy degradation similarly accumulates immune activated cells (Fig 1D) as do other cargo adapters; BNIP3L, OPTN, FUNDC1 and TAX1BP1 (mitophagy) and FAM134B, SEC62, ATL3 and TEX64 (ER-phagy) (Fig 1C, S1B). The accumulation of autophagy machinery is a relatively rapid response detected within 3-6 hours of antigen receptor triggering (Fig 1E). Notably, increases in the abundance of proteins that mediate autophagy were not correlated with increased abundance of their mRNA (Fig 1A, 1D, S1A, S1B). Indeed, *Map1lc3b*, *Gabarapl2* and *Sqstm1* mRNA decreased in antigen activated CD8 T-cells (Fig 1A, 1D, S1A, S1B).

**Figure 1:**
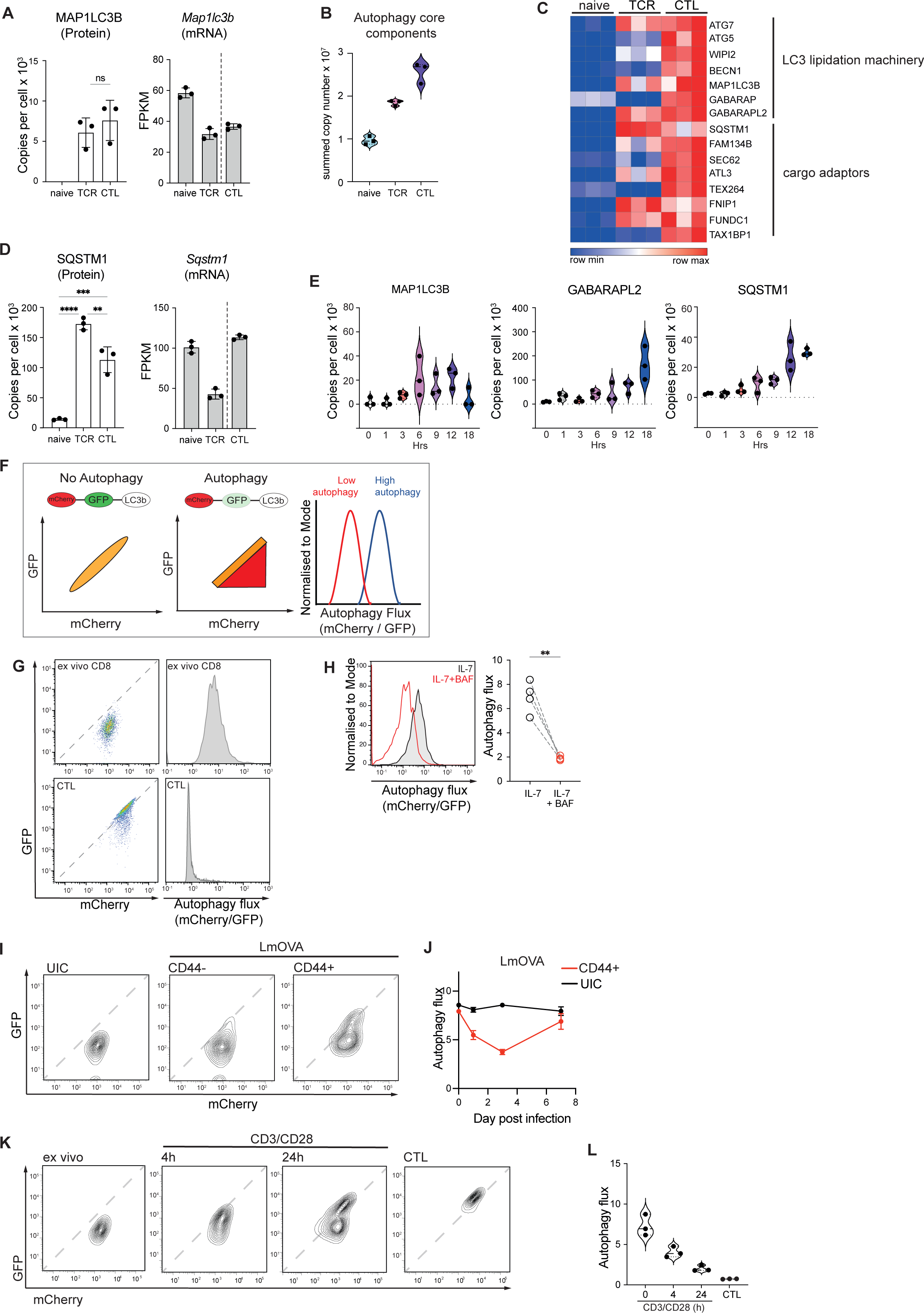
Immune activated T cells repress autophagy flux. **A** Quantitative proteomics data showing mean protein copy number per cell (left) and RNASeq data shown as FPKM (right) from naïve, 24 hour TCR activated P14 CD8 T cells and IL-2 maintained effector CD8 T cells (CTL) for MAP1LC3B**. B** Summed copy numbers of core autophagy components from naïve, 24 hour TCR activated P14 CD8 T cells and CTL, based on list curated by Bordi et al ^63^. The list of proteins included is provided in Supplementary table S1. **C** Heat map of autophagy machinery expression during CD8 differentiation (naïve, 24 hour TCR activated P14 CD8 T cells and IL-2 maintained CTL). **D** Protein copy numbers (left) and RNASeq data (right) from naïve, TCR activated and CTL for SQSTM1. **E** Mean protein copy numbers from naïve, and OT1 CD8 T cells activated with SIINFEKL for the indicated times of MAP1LC3B, GABARAPL2 and SQSTM1. **F** Schematic representations of flow cytometry data generated with cells from mCherry:GFP:LC3b autophagy reporter mice: scatter plot showing equivalent fluorescence of GFP and mCherry indicating no autophagy (left) and scatter plot showing a loss of GFP fluorescence with a maintenance of mCherry fluorescence indicating ongoing autophagy (centre). A derived parameter of mCherry/GFP fluorescence can be plotted as a histogram, and the mean fluorescence intensity (MFI) calculated. A value of 1 is indicative of low autophagy; a shift of mCherry/GFP above 1 is indicative of ongoing autophagy flux (right). **G** Representative flow cytometry data showing GFP:mCherry fluorescence profiles of ex vivo, naïve CD8 cells and IL-2 maintained effector CD8 T cells (CTL) (left) and representative histograms showing Autophagy flux derived parameter (mCherry/GFP; right). **H** Representative flow cytometry data showing Autophagy flux in IL-7 maintained naïve CD8 T cells with or without Bafilomycin A (BAF, 200nM) for 5 hrs (left) the mean fluorescence intensity (MFI) are shown in the right panel. **I, J** GFP:mCherry:LC3b autophagy reporter mice were infected with attenuated OVA expressing *Listeria monocytogenes* (LmOVA). Splenic CD8 T cells were analysed on 1, 3 and 7 days post-infection. **I** Representative flow cytometry data showing GFP:mCherry fluorescence profiles of CD8 cells from uninfected control (UIC) and LmOVA infected CD44- and infected CD44+ T cells 3 days post infection. **J** Autophagy Flux (mCherry/GFP) of CD44+ CD8 T cells from LmOVA infected mice, or CD8 T cells from uninfected controls over the extended timecourse. **K,L** Flow cytometry scatter plots (**K**) and autophagy flux (**L**) showing GFP:mCherry fluorescence profiles from autophagy reporter CD8 T cells: *Ex vivo*, 4 and 24 hours after TCR activation (anti-CD3/CD28) and in IL-2 maintained effector cells (CTL). Proteomic data (A-D) are from Howden et al ^64^, all proteomic data (A-E) are available on Immpres.co.uk ^65^. FPKM mRNA data from Spinelli et al ^66^ (A,D). All data are from a minimum of 3 biological replicates. Data points in bar charts are indicative of biological replicates. Error bars represent the mean +/- SD. P ≤ 0.0001 indicated by ****; P ≤ 0.0005 = ***; P ≤ 0.001 = **; P ≤ 0.05 = *

If cells have high levels of autophagic flux, there will be constant degradation of ATG8 proteins and cargo adaptors such as p62/SQSTM1. Consequently, these proteins would be low abundance when cells have high autophagy flux but would accumulate under conditions of low autophagy. The low abundance of cargo adaptor proteins, MAP1LC3b and the GABARAPs in naïve T cells and their accumulation in antigen activated CD8 T cells could thus reflect that naive T cells have high levels of autophagy but respond to antigen by rapidly repressing autophagy and sustain a suppressed state of autophagy as they differentiate to effector cells. To explore this hypothesis, we used an autophagy flux reporter mouse model that incorporates normalization of autophagosome levels to allow more accurate measurements of autophagy flux in primary cells and tissues ^21,22^. In this model, cDNA encoding an mCherry:GFP:Map1lc3b (mCherry:GFP:LC3b) fusion protein is engineered into the ROSA 26 locus of a WT (C57 B6/J) mouse to allow ubiquitous expression. When autophagy initiates, the tandem-tagged LC3b is recruited into autophagosomes and when these fuse with lysosomes, the low luminal pH of the resulting autolysosome quenches the GFP fluorophore. mCherry fluorescence emissions are insensitive to these pH changes and hence increased autophagic flux can be quantified by comparing the magnitude of GFP fluorescence quenching normalized to the mCherry signal. Scatter plots comparing GFP and mCherry fluorescence give a visual indication of populational autophagy as there will be a linear relationship between GFP and mCherry fluorescence if cells are not undergoing autophagy. If the LC3b fusion protein is recruited to an autophagosome the GFP fluorescence will be quenched, and cells will display a lower GFP than mCherry signal. A measure of autophagy flux can thus be calculated by dividing the mCherry signal by the GFP signal (Figure 1F).

The data show that both *ex vivo* naive CD8 T-cells and in vitro generated cytotoxic T-cells from LC3b reporter mice have a strong mCherry signal, indicative of LC3b reporter expression (Fig 1G, supp 1C). However, there is clear quenching of GFP fluorescence signal in naïve CD8 T-cells, whereas there was a high linear correlation of the GFP and mCherry signal in CTL (Fig 1G). The quenching of the GFP signal in the naive T cells indicates that the autophagy reporter fusion protein is in an acidic compartment in these cells but not in the CTL. Bafilomycin A1, which inhibits lysosomal V-ATPase activity, blocks the final step of autophagy by raising lysosomal pH and preventing the fusion of lysosomes and autophagosomes ^23,24^. Fig 1H shows that the high levels of autophagy flux seen in naïve T cells are reduced when cells treated with Bafilomycin A1. We also examined autophagy flux in immune activated CD8 T cells *in vivo* in a model where mCherry:GFP:LC3b reporter mice were immunised with a modified strain of OVA expressing *Listeria monocytogenes* (LmOVA). The data show that CD8 T cells activated in response to Listeria infection have reduced autophagic flux early after the immune challenge/infection, D1 and D3, but that autophagy levels return to those seen in control uninfected mice by day 7 (Fig 1I, J). Collectively these analyses of quiescent and immune activated T cells *in vitro* and *in vivo* are consistent with the model of high levels of autophagic flux in naive T cells versus low levels of autophagic flux in immune activated effector CD8 T cells.

In further work we examined how rapidly T cell activation impacted autophagy flux. In these experiments, CD3/CD28 antibodies were used to polyclonally stimulate naïve CD8+ T cells expressing the mCherry:GFP:LC3b reporter and the GFP:mCherry fluorescence profiles of the cells was analysed over time. The data (Fig 1K and 1L) show a populational shift with the emergence of cells with a non GFP quenched signal in the autophagy reporter. After 24 hours of CD3/CD28 stimulation, a large proportion (∼70%) of the cells were now linear for GFP:mCherry fluorescence (Fig 1K and 1L). The ability to detect reductions in autophagic flux with a few hours of antigen receptor engagement is consistent with the timing of the accumulation of ATG8 family members and cargo adaptors in antigen activated CD8 T cells (Fig 1E).

### Pro-inflammatory cytokines repress autophagy flux

Figure 1 shows that CTL generated by culturing antigen activated CD8 T cells in IL-2 have linear GFP:mCherry fluorescence of an autophagy reporter consistent with low autophagy flux in these cells. To address the role of IL-2 in repressing autophagy in CTL, we monitored the impact of cytokine deprivation on the autophagy reporter. The data (Fig 2A, 2B) show that when CTL are IL-2 deprived, or if CTL are sustained in IL-2 but treated with inhibitors of Jak tyrosine kinases to block IL-2 signalling, they revert to a profile of quenched GFP: mCherry fluorescence, ie high autophagy flux. The suppression of autophagy flux in CTL is thus dependent on sustained IL-2 signalling. Consistent with increased autophagy flux there was decreased protein expression of ATG8 family members and autophagy adaptor levels in IL-2 deprived CTL but not decreased mRNA (Fig 2C, S2A). In further experiments we examined the ability of other cytokines to regulate autophagy flux in antigen primed CTL. We focused on the impact of the cytokines IL-12 and IL-18 which drive terminal effector CD8 T cell differentiation and IL-15 which promotes differentiation of memory T cells. The data show that autophagy flux is repressed (Fig 2D) and ATG8 family members and autophagy cargo adaptors accumulate in IL-12+IL18 maintained CTL (Fig 2E, S2B). In contrast, antigen activated CD8 T cells expanded with IL-15 do not suppress autophagy flux (Fig 2F, 2G). Consistent with high autophagy flux in IL-15 maintained CD8-T cells there is also decreased expression of ATG8 family members and cargo adaptors in IL15 maintained CD8 T cells at the protein but not mRNA level (Fig 2H, S2C). Collectively these data argue that naïve and memory T cells have high levels of autophagic flux but when naïve cells respond to antigen receptor engagement autophagic flux is repressed. Pro-inflammatory cytokines such as IL-2 and IL-12 that drive terminal effector T cell differentiation repress autophagy in antigen primed CD8 T-cells whereas cytokines that support memory CD8 T-cell homeostasis such as IL-15 do not.

**Figure 2:**
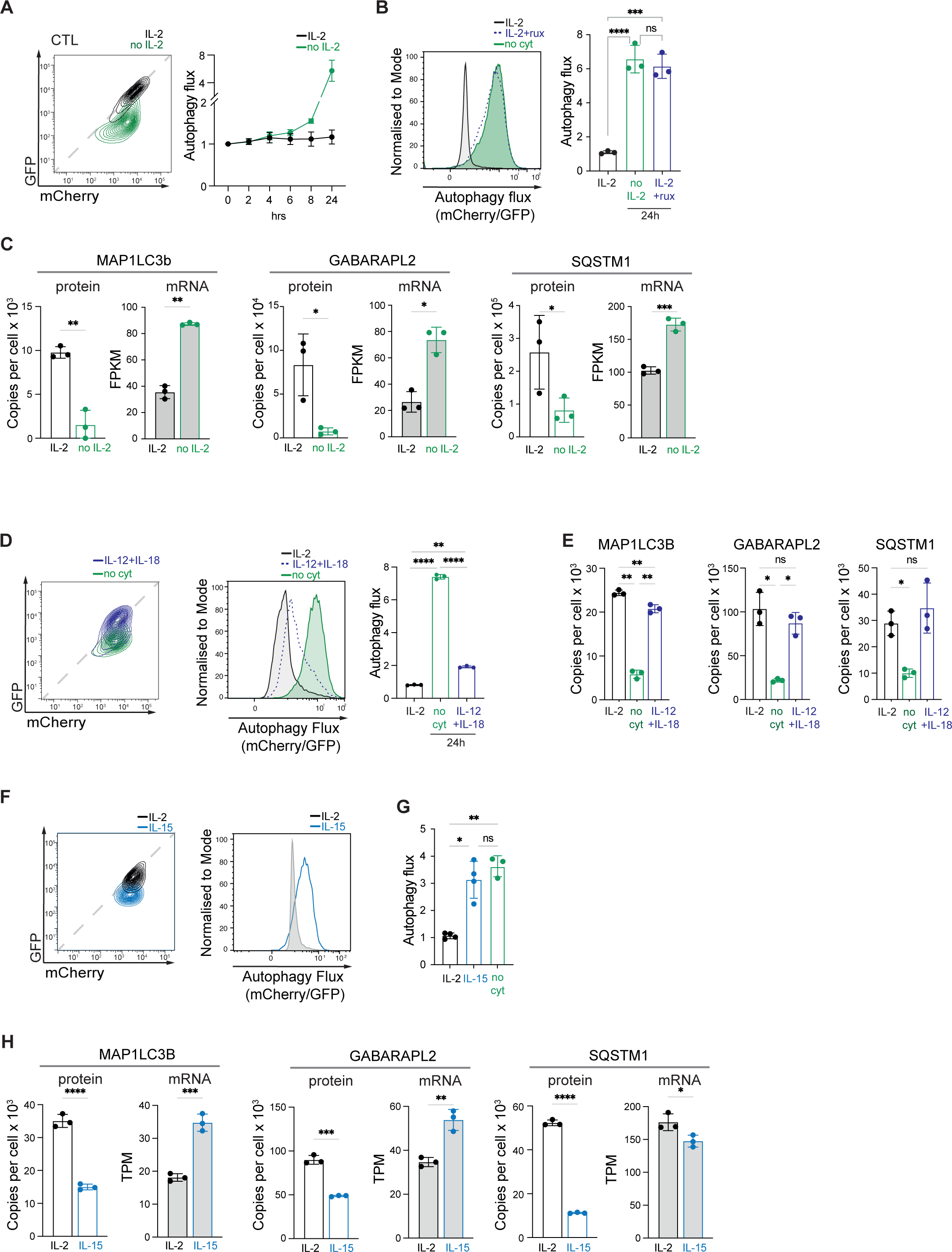
Pro-inflammatory cytokines repress autophagy flux. **A** Representative GFP:mCherry fluorescence profiles from autophagy reporter CTL maintained with or without IL-2 over 24h (left). The right panel shows the Autophagy flux (mCherry/GFP) values over the expanded time course. **B** Representative flow cytometry data showing Autophagy flux (mCherry/GFP) from autophagy reporter CTL maintained with or without IL-2 or the JAK inhibitor ruxolitinib (1uM) for 24hr (left). The graph (right) shows the mean Autophagy flux (mCherry/GFP) values. **C** Mean protein copy number per cell and RNASeq data (shown as FPKM) from CTL maintained with or without IL-2 over 24h for MAP1LC3B, GABARAP2L and SQSTM1. **D** GFP:mCherry fluorescence profiles (left), autophagy flux histogram (centre) the mean Autophagy flux (mCherry/GFP) values (right) from autophagy reporter CTL maintained with IL-2, with IL-12+IL-18 or no cyokine for 24h. **E** Mean protein copy number per cell from CTL maintained with IL-2, with IL-12+IL-18 or no cytokine for 24h for MAP1LC3B, GABARAP2L and SQSTM1. **F** GFP:mCherry fluorescence profiles (left) and autophagy flux histogram (right) from autophagy reporter CD8 T cells expanded with IL-2 (CTL) or expanded with IL-15 (memory-like). Mean autophagy flux (mCherry/GFP) values from autophagy reporter IL-2 (CTL), IL-15 (memory-like) or deprived of cytokine for 24hr. **H** Mean protein copy number per cell and RNASeq data (shown as TPM) from CD8 T cells expanded with IL-2 (CTL) or expanded with IL-15 (memory-like) for MAP1LC3B, GABARAP2L and SQSTM1. The data in 2C are from Spinelli et al ^66^; 2E are available on impress.co.uk ^65^. 2H are from Marchingo et al ^67^. Data are from a minimum of 3 biological replicates. Error bars represent the mean +/- SD. Data points in bar charts are indicative of biological replicates. Data points in time courses represent the mean +/- SD. P ≤ 0.0001 indicated by ****; P ≤ 0.0005 = ***; P ≤ 0.001 = **; P ≤ 0.05 = *

### Amino acid supply controls autophagy in CD8 T cells

One key autophagy stimulus is amino acid deprivation ^25^. In this respect, naive T cells and IL-15 maintained memory CD8 T cells have low levels of amino acid transport compared to high levels of amino acid transport in antigen activated T cells and IL-2 maintained CTL ^26,27^, reflective of lower amino acid transporter expression (Supp S3A,B,C). Moreover, comparisons of the timing of autophagy repression versus amino acid transporter upregulation shows a clear inverse relationship between autophagy flux and amino acid transporter expression in antigen receptor activated CD8+ T cells (Fig 1L vs Fig 3A). CD8 T cells activated for 24hr with CD3/CD28 antibodies show heterogeneity of autophagic flux (Fig 1J, Fig 3B). The majority of cells showed linear GFP:mCherry fluorescence indicating repressed autophagy, however there was a substantial subset of cells that retained quenched GFP fluorescence (Fig 1J, Fig 3B). Accordingly, we addressed whether the heterogeneity in autophagy repression was linked to any heterogeneity in amino acid transport in this polyclonal population. Fig 3B shows that the subpopulation of cells with high levels of autophagic flux (ie low GFP fluorescence and high mCherry fluorescence), have low amino acid transport capacity whereas the cells that have repressed autophagy have high amino acid transport capacity (Fig 3B). We also observed that autophagy repression in CD8 T cells activated *in vivo* correlates with their amino acid transport capacity. Fig 3C shows amino acid uptake capacity and autophagy flux of activated CD8 T cells from mice 3 days and 7 days after Listeria infection. On D3, the activated CD8 T cells that have high amino acid transport capacity also show low autophagy flux. Activated CD8 T cells analysed on D7 post listeria infection have low amino acid transport capacity and high autophagy flux, similar to non-infected controls (Fig 3D). This is consistent with a model whereby the mechanism that underpins the ability of activated T cells to switch off autophagy is linked with the upregulation of amino acid transport and the intracellular sensing of amino acids imported from the environment.

**Figure 3:**
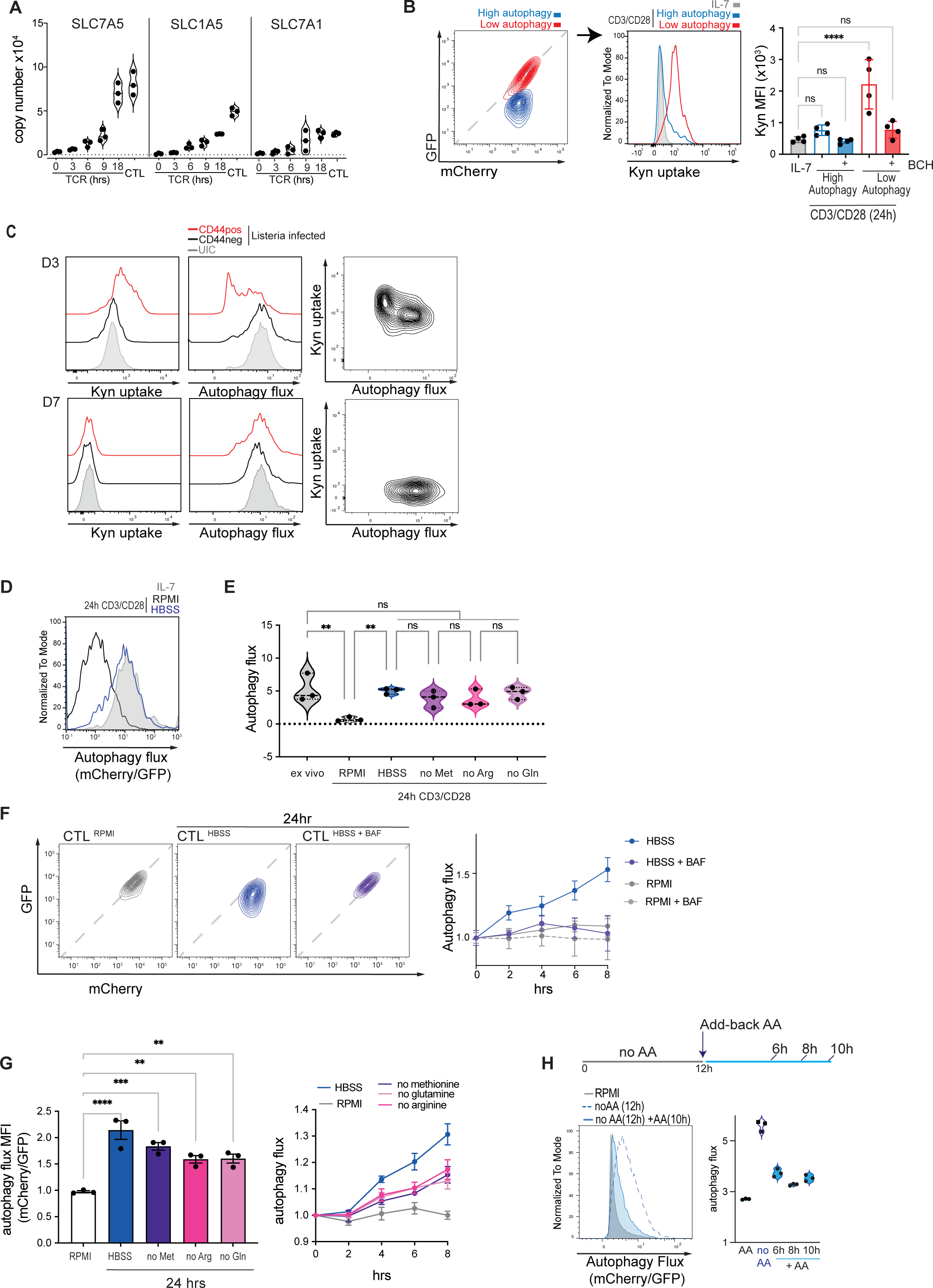
Amino acid supply controls autophagy in CD8 T cells. **A** Mean protein copy numbers of SLC7A5 (left), SLC1A5 (center) and SLC7A1 (right) from CTL and OT-1 CD8 T cells activated with SIINFEKL for the indicated times. **B** The data show representative flow cytometry GFP:mCherry fluorescence of CD8 T cells from autophagy reporter mice after 24h CD3/CD28 stimulation (left). Cells undergoing high or low autophagy flux are indicated. The centre panel shows a representative histogram of kynurenine (Kyn) uptake in IL-7 maintained or CD3/CD28 activated CD8 cells undergoing high or low autophagy. The right panel shows the associated mean fluorescence intensity (MFI) of kynurenine uptake in the presence or absence of the System L competitive substrate, BCH. **C** Representative histograms showing kynurenine uptake (left panels) and Autophagy flux (mCherry/GFP) profiles of autophagy reporter CD8 cells from uninfected control (UIC) and LmOVA infected CD44- and infected CD44+ T cells 3 (top) and 7 (bottom) days post infection. The scatter plots show kynurenine uptake against autophagy flux (right panels). **D** Representative Autophagy flux (mCherry/GFP) profiles of autophagy reporter CD8 cells maintained in IL-7 or activated through the TCR (CD3/CD28) for 24hr in the presence (RPMI) or absence of amino acids (HBSS). **E** Mean Autophagy flux (mCherry/GFP) values from autophagy reporter CD8 cells either *ex vivo* or activated through the TCR (CD3/CD28) for 24hr in the presence (RPMI) or absence of amino acids (HBSS) or in RPMI lacking individual amino acids; methionine (no Met), arginine (no Arg) or glutamine (no Gln). **F** Representative flow cytometry data showing GFP:mCherry fluorescence profiles of CTL maintained in RPMI, or switched into amino acid free HBSS with or without Bafilomycin A (BAF;200nM) for 24hr. The graph (right) shows mean Autophagy flux (mCherry/GFP) values for these conditions over 0-8hrs. **G** Mean Autophagy flux (mCherry/GFP) values from autophagy reporter CTL maintained in RPMI, or switched into amino acid free HBSS or RPMI lacking individual amino acids; methionine (no Met), arginine (no Arg) or glutamine (no Gln) for 24hr. The graph (right) shows mean Autophagy flux (mCherry/GFP) values for these conditions over 0-8hrs. **H** Representative Autophagy flux (mCherry/GFP) profiles of autophagy reporter CTL maintained in the presence of full amino acids (RPMI), depleted of amino acids (noAA; in HBSS) for 12 hrs, or depleted of amino acids for 12hrs and then switched into full amino acids (RPMI) for 10 h (left histogram). The graph (right) shows mean autophagy flux (mCherry/GFP) values from autophagy reporter CTL maintained in the presence of full amino acids (RPMI), depleted of amino acids (noAA; in HBSS) for 12 hrs or depleted of amino acids for 12hrs and then switched into full amino acids (+AA, RPMI) for 6h, 8h, or 10 h. Proteomic data (A) are from Immpres.co.uk. Data are from a minimum of 3 biological replicates. Data points in bar charts are indicative of biological replicates. Error bars represent the mean +/- SD. P ≤ 0.0001 indicated by ****; P ≤ 0.0005 = ***; P ≤ 0.001 = **; P ≤ 0.05 = *

To test a causal link between amino acid supply and autophagy we examined whether immune activated CD8 T cells could repress autophagy if they were amino acid deprived. CD8 T cells expressing the mCherry:GFP:LC3b reporter were accordingly activated in amino acid free media HBSS or in the absence of the single amino acids, glutamine, arginine or methionine. The data show that immune activated T cells cannot repress autophagy unless they have an extracellular amino acid supply (Fig 3D); deprivation of a single amino acid also prevents autophagy repression in response to immune activation (Fig 3E). The data also show that if IL-2 maintained CTL are switched from RPMI, an amino acid replete media, to the amino acid free HBSS they rapidly quench the GFP fluorescence in the mCherry:GFP:LC3b reporter consistent with increased autophagic flux (Fig 3F). When amino acid deprived CTL are treated with Bafilomycin A1 there is maintenance of GFP fluorescence and linear mCherry:GFP fluorescence of the LC3b reporter (Fig 3F). Strikingly, even depletion of single amino acids such as glutamine, arginine or methionine was sufficient to induce autophagy in CTL (Fig 3G). Importantly, re-addition of amino acids to amino acid deprived CTL re-establishes autophagy repression (Fig 3H). Hence, the suppression of autophagic flux in CTL is dynamic and dependent on sustained amino acid supply.

### CTL control autophagy by VPS34 dependent and AMPK and mTORC1 independent pathways

The class III Phosphatidylinositol 3 Kinase, Vacuolar protein sorting 34 (VPS34; PIK3C3) is critical for autophagosome formation ^28–30^ and has been shown to control autophagy in T cells ^5,6,28,31^. In this context, autophagy flux in amino acid deprived CTL is dependent on VPS34 activity, as demonstrated by using the highly specific VPS34 inhibitor, VPS34-IN1 ^32^(Fig 4A). Furthermore, the data also show that the high levels of autophagy flux present in IL-7 maintained naïve T cells are dependent on VPS34 (Fig 4B).

**Figure 4:**
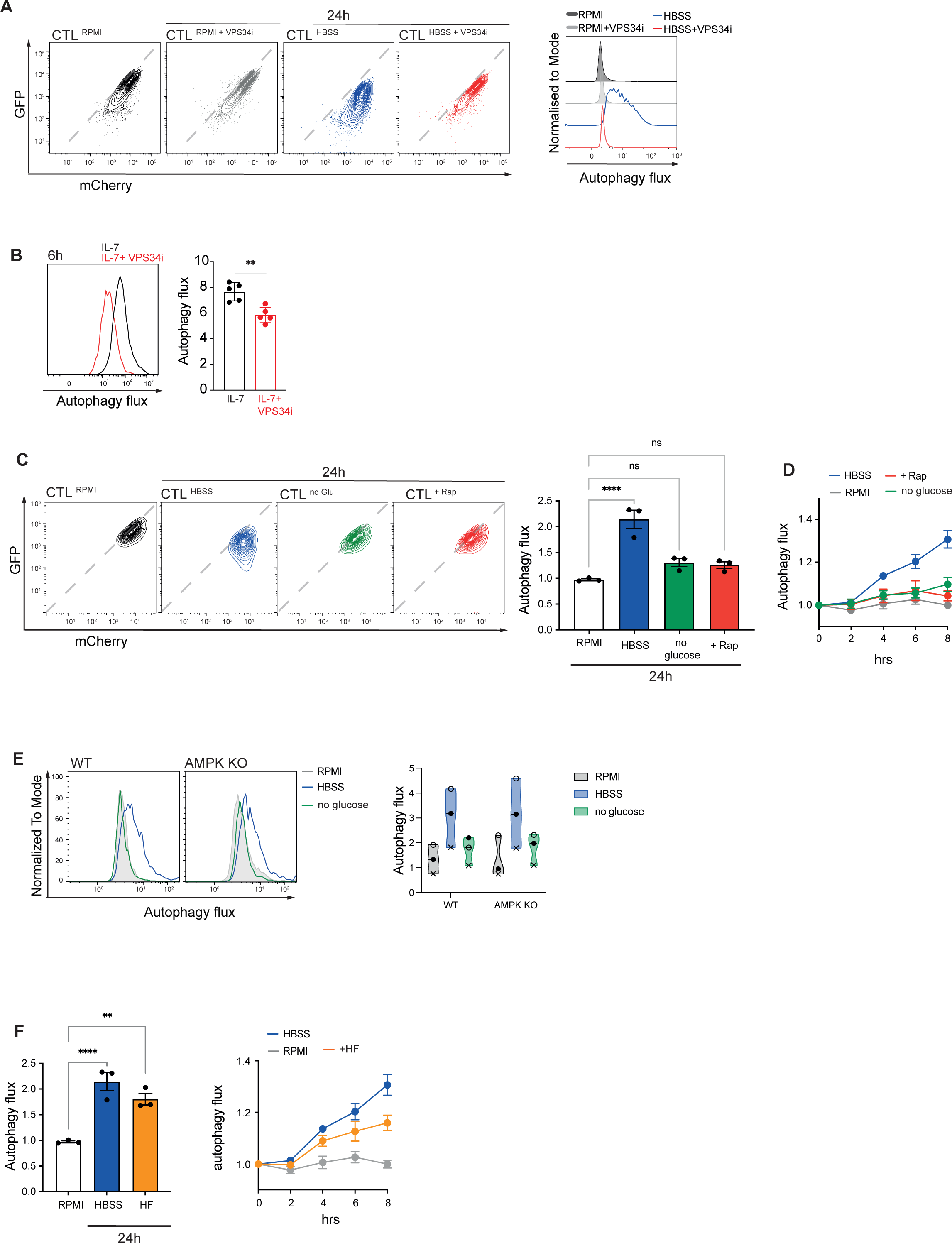
VPS34 dependent and AMPK and mTORC1 independent autophagy in CD8 T cells. **A** Representative mCherry:GFP flow cytometry plots of autophagy reporter CTL maintained in the presence of full amino acids (RPMI) with or without VPS34 inhibitor (VPS34i) or depleted of amino acids (noAA; in HBSS) with or without VPS34i for 24h (left panels). The graph (right) shows the corresponding Autophagy flux (mCherry/GFP) histogram. **B** Autophagy flux histogram (left) and the mean Autophagy flux (right) of IL-7 maintained naïve CD8 T cells with or without VPS34i for 6h. **C** Representative mCherry:GFP plots (left) and corresponding mean autophagy flux (mCherry/GFP) values (right) of autophagy reporter CTL maintained in the presence of full amino acids (RPMI), depleted of amino acids (noAA; in HBSS), switched into glucose free RPMI (noGlu) or treated with rapamycin (Rap, 20nM) for 24h. **D** The autophagy flux (mCherry/GFP) values over the expanded time course of CTL treated as in **C**. **E** Representative autophagy flux (mCherry/GFP) histograms (left) and mean Autophagy flux (mCherry/GFP) values (right) of CTL from control (WT) autophagy reporter mice or AMPK KO autophagy reporter mice maintained in the presence of full amino acids (RPMI), depleted of amino acids (noAA; HBSS) or switched into glucose free RPMI (noGlu) for 24 hrs. **F** Mean autophagy flux (mCherry/GFP) values of autophagy reporter CTL maintained in the presence of full amino acids (RPMI), depleted of amino acids (noAA; HBSS) or treated with Halofuginone (HF; 100nM) for 24h (left panel) and over the expanded time course (right panel). Data points in bar charts are indicative of biological replicates. Error bars represent the mean +/- SD. Different experiments are indicated by different data points in the violin plot (**E**). P ≤ 0.0001 indicated by ****; P ≤ 0.0005 = ***; P ≤ 0.001 = **; P ≤ 0.05 = *

One protein kinase reported to regulate autophagy is the AMP activated protein kinase (AMPK) ^33^. The activation of AMPK in CD8 T cells is required for memory T cell differentiation and occurs under conditions of glucose deprivation ^34,35^. Moreover, one role for AMPK in CTL is to repress activation of mTORC1 (mammalian target of rapamycin complex 1) when glucose is limited ^34^. In this context, AMPK and mTORC1 have been shown to regulate autophagy in a number of cell systems, both in positive and negative manner ^33,36–38^. We therefore assessed if glucose deprivation or inhibition of mTORC1 would increase autophagic flux in CTL. The data show that while amino acid deprivation suppresses mTORC1 activity (Fig S4A), only amino acid deprivation, not direct mTORC1 inhibition, induces autophagic flux in CTL (Fig 4C, D). Moreover, glucose deprivation, which activates AMPK and inhibits mTORC1 (Supp Fig S4A, S4B), is not associated with induction of autophagic flux (Fig 4 C, D). We also explored autophagy regulation in in AMPK α1 null T cells and found no significant difference in basal levels of autophagy in WT and AMPK KO CTL (Fig 4E). Likewise, acute removal of amino acids but not glucose induced autophagy in both WT and AMPK KO CTL (Fig 4E). One other candidate to regulate autophagy in response to amino acid deprivation is the serine threonine kinase GCN2, which is activated by the uncharged tRNAs that accumulate under amino acid starvation ^39–43^. Halofuginone, HF, acts as a GCN2 agonist by inhibiting prolyl-tRNA synthetase and thereby mimicking the unavailability of proline and has been shown to induce amino acid starvation responses in T cells ^44–47)^. The present data show that this GCN2 agonist induces autophagic flux in CTL (Fig 4F).

### VPS34 control of autophagy and protein degradation in cytotoxic T cells

The ability of VPS34 inhibition to supress autophagy in CTL opens a strategy to explore how autophagy shapes autophagic CTL proteomes. We thus used mass spectrometry to quantitively analyse CTL proteomes to identify proteins whose expression is regulated by VPS34 dependent pathways in both nutrient rich conditions (low autophagy) or amino acid deprived conditions, which drives high autophagy (Fig 5A). The rationale is that in CTL undergoing autophagy, VPS34 inhibition would supress autophagy and increase expression of proteins degraded via VPS34 dependent signalling pathways. Conversely, proteins fuelled by autophagy would decrease in expression in autophagic T cells treated with VPS34 inhibitors because the inhibition of autophagy would prevent autophagy dependent recycling of amino acids for protein synthesis.

**Figure 5:**
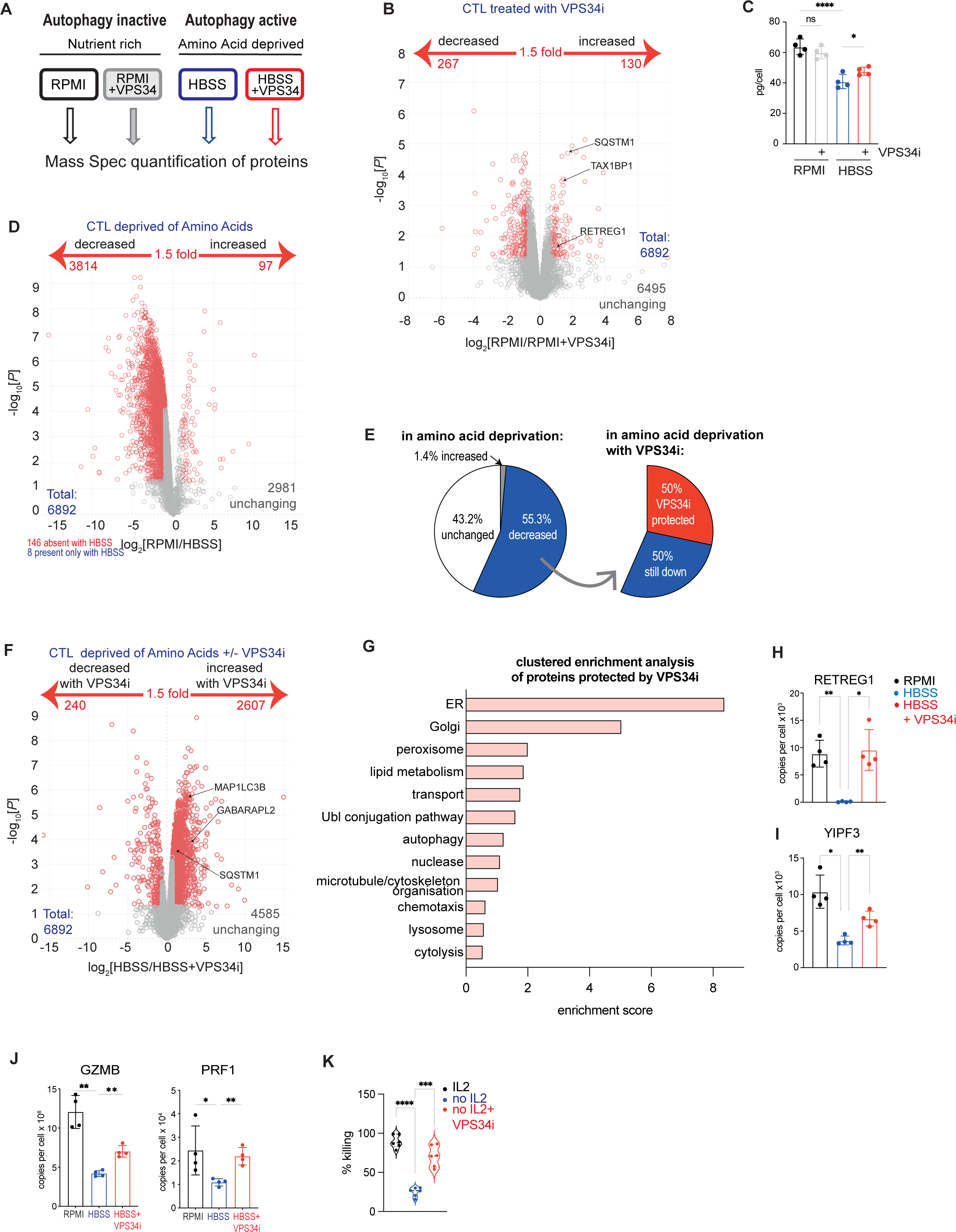
VPS34 control of autophagy and protein degradation in CTL during AA starvation. **A** Schematic diagram showing the 4 CTL treatment conditions (18hr) for subsequent proteomic analysis. The full list of proteins detected is available in Supplementary Table S5. **B** Volcano plot showing the ratio changes for proteins expressed in CTL in RPMI with or without VPS34i treatment. *P* values below 0.05 and fold change above 1.5 are considered significant and marked in red. **C** Total protein content calculated from proteomics of CTL in amino acid replete media (RPMI) or amino acid deprived media (HBSS) with or without VPS34i for 18hr. **D** Volcano plot showing the ratio changes for proteins in CTL in RPMI compared with amino acid deprived (HBSS) CTL. *P* values below 0.05 and fold change above 1.5 are considered significant and marked in red. **E** The proportion of the CTL proteome changed upon amino acid starvation (RPMI into HBSS; left) and the proportion of the amino acid starved CTL proteome which was increased upon VPS34 inhibition (HBSS to VPS34i; right). **F** Volcano plot showing the ratio changes of proteins in amino acid deprived (HBSS) CTL compared to amino acid deprived (HBSS) CTL treated with VPS34i. *P* values below 0.05 and fold change above 1.5 are considered significant and marked in red. **G** Clustered enrichment analysis on proteins from amino acid deprived (HBSS) CTL that were significantly increased with VPS34i treatment. The full enrichment table is available in Supplementary data file S5. **H-J** Quantitative proteomics data showing mean protein copy numbers per cell of RETREG1 (H), YIPF3 (I), GZMB and perforin (PRF1) (J). **K** Target cell killing capacity of CTL maintained in IL-2 or deprived of IL-2 with or without VPS34i for 24h. Data points in bar charts are indicative of biological replicates. Error bars represent the mean +/- SD. P ≤ 0.0001 indicated by ****; P ≤ 0.0005 = ***; P ≤ 0.001 = **; P ≤ 0.05 = *

We first investigated the effect of VPS34 inhibition on the proteomes of CTL cultured in amino acid replete medium, where autophagy flux is low. In these experiments 6892 proteins were quantified and expression of only 130 proteins increased following VPS34 inhibition (Fig 5B and Supp file S5). These included many autophagy related proteins such as TAX1BP1, p62/SQSTM1, RETREG1, MAP1LC3B and GABARAPL2 (Fig 5B, S5A, Fig S5B; supplemental tableS5). The failure to see any significant impact of VPS34 inhibition on CTL proteomes when cells are not doing autophagy highlights how VPS34 dependent pathways are not needed for IL-2 signalling or for the control of protein synthesis and cell metabolism under nutrient replete conditions.

We then looked the impact of VPS34 inhibition on CTL proteomes in cells with high autophagic flux because of amino acid deprivation. In the absence of amino acids for 18 hours, CTL decrease expression of approximately 3800 proteins, and lose approx. 1/3 of cell mass (Fig 5C, D). The proteins that are downregulated under these conditions will include proteins that are degraded by the canonical proteosome degradation pathway but then not resynthesized due to the unavailability of amino acids in this model system. There will, however, be some proteins whose expression is decreased because of the increases in autophagic flux that occur in amino acid deprived CTL. This latter group of proteins can be identified by their failure to be degraded when the VPS34 inhibitor is used to prevent autophagy. Here the striking result was that VPS34 inhibition repressed the degradation of <2000 proteins in amino acid deprived T cells ie approximately 50% of the proteins downregulated in amino acid deprived CTL were degraded by VPS34 sensitive mechanisms (Fig 5E, 5F). The loss of cell mass in amino acid deprived CTL was thus blunted when autophagy was inhibited by the VPS34 inhibitor (Fig 5C).

Proteins ‘protected’ from degradation in the amino acid deprived VPS34 inhibitor treated CTL included components of the autophagy machinery such as SQSTM1, GABARAPL2 and MAP1LC3B (Fig 5F, S5B). Clustered enrichment analysis indicated that one dominant effect of VPS34 inhibition on amino acid deprived CTL was to retain expression of ER and Golgi related proteins (Fig 5G and Supp Fig S5C, table S5). In this respect inhibition of autophagy in amino-acid deprived CTL results in the accumulation of the ER-phagy and Golgi-phagy receptors RETREG1 and YIPF3 (Fig 5H and 5I). The enrichment analysis also highlighted that proteins involved in controlling and mediating T cell cytotoxic function were retained with VPS34 inhibition including RAB27A, RAB27B, STXBP2, STXBP3 and UNC13D (Fig 5G and Supp Fig S5D, S5E).

In particular, a striking observation was that CTL deprived of amino acids downregulate expression of effector cytolytic proteins, such as perforin and granzymes (Fig 5J, Supp Fig S5D). The full loss of these cytolytic effector molecules was prevented by VPS34 inhibition of autophagy (Fig 5J, Supp Fig S5E). Furthermore, the reduced killing capacity of effector IL-2 deprived CTL, which drives high autophagy flux (Fig 2A), is restored upon VPS34 inhibition (Fig 5K). Collectively these data reveal that granzymes and perforin are autophagy substrates in T cells and repression of autophagy is required to sustain their expression and to sustain CTL killing capacity.

Finally, we noted that amino acid deprived autophagic CTL showed selective remodelling of mitochondria (Fig 6A). Some of this remodelling was prevented when autophagy was inhibited, and some was not. For example, mitochondrial proteins involved in the electron transport chain were downregulated by amino acid starvation, but these losses were not prevented by autophagy inhibition (Fig 6B and S6A). In contrast, expression of the mitochondrial pyruvate carrier MPC2 (Fig 6C), TCA cycle proteins (Fig 6D), and key proteins involved in Fatty acid oxidation (Fig 6E) and mitochondrial amino acid metabolism (Fig 6F) were decreased upon amino acid deprivation and this was prevented by VPS34 inhibition. One other important category of proteins downregulated in amino acid starved T cells but protected by VPS34 inhibition of autophagy were nutrient transporters notably, the amino acid transporters SLC1A5, SLC7A1, SLC7A5 and SLC7A6 (Fig 6G); glucose transporters SLC2A1, SLC2A3 as well as lactate transporters, SLC16A1 and SLC16A3 (Fig 6H and 6I). Cellular levels of glucose transporters are rate limiting for T cell activation as glucose is a key fuel to produce ATP from oxidative phosphorylation and glycolysis ^48,49^). As well, the expression of lactate transporters is rate limiting for glycolysis ^49^. Collectively these data predict that amino acid deprived T cells will downregulate glycolysis, but inhibition of autophagy will prevent this. The data also predict reductions in oxidative phosphorylation in amino acid deprived CTL but the failure of autophagy inhibition to prevent the loss of major oxidative phosphorylation components (Fig 6B and Supp Fig S6A) predicts that these changes would not be prevented by autophagy inhibition.

**Figure 6:**
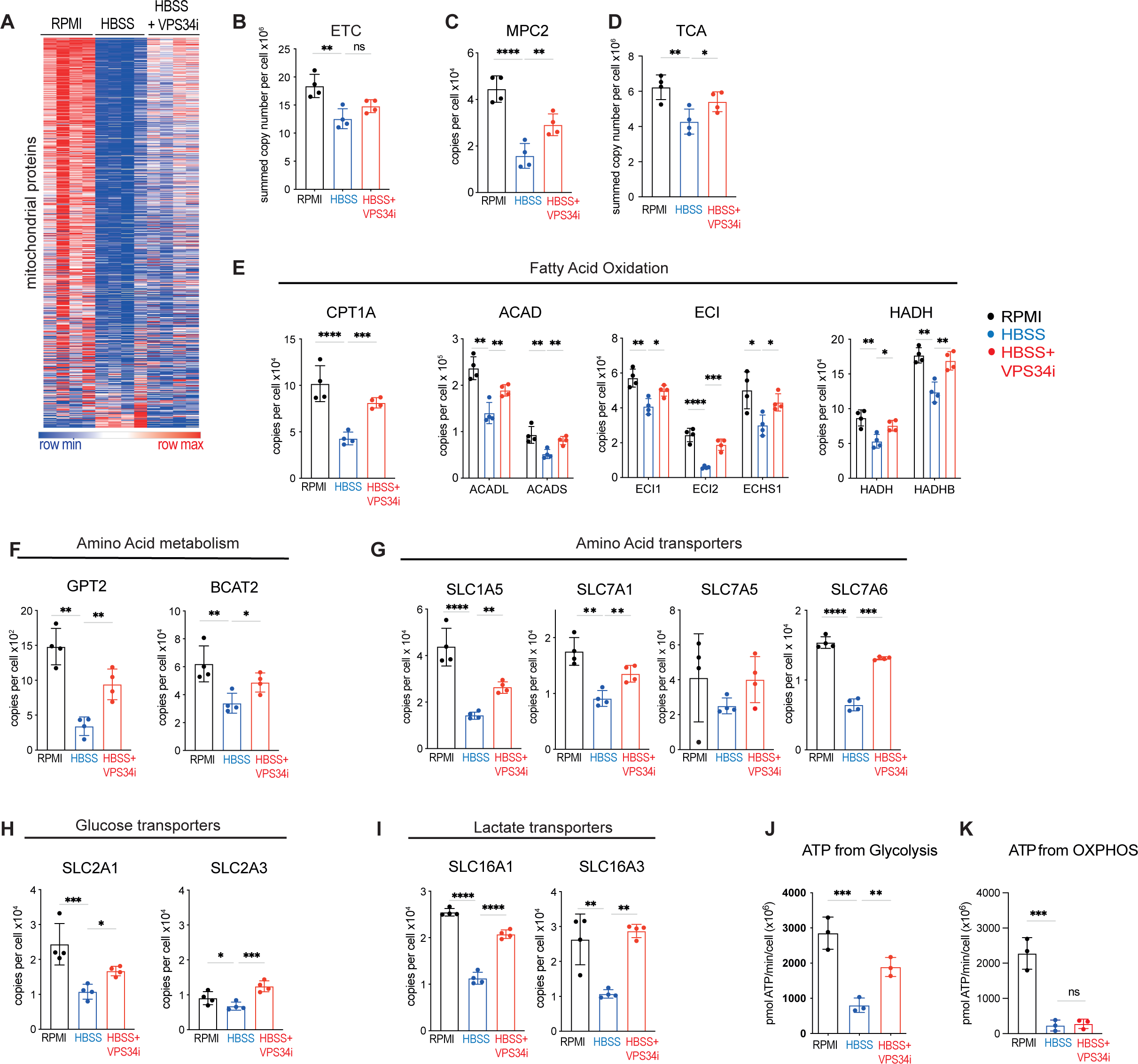
CTL metabolic remodelling during AA starvation is restrained by VPS34 inhibition. **A** Heatmap of mitochondrial proteins (GO 0005739, MitoCarta3.0) from CTL in replete amino acids (RPMI), or amino acid deprived (HBSS) with or without VPS34i for 18hr. The full list of proteins within the heatmap is provided in Supplementary data file S6. **B** Summed protein copy number of electron transport chain (ETC; MitoCarta3.0 ‘OXPHOS’) **C** Protein copy numbers per cell of MPC2. **D** Summed protein copy number of TCA cycle proteins (KEGG M00009) **E-I**Protein copy numbers per cell of enzymes and regulators involved in Fatty Acid Oxidation (E); Amino Acid metabolism (F); Amino Acid transporter (G); Glucose transporter (H) and Lactate transporter (I) as indicated. **J,K** Predicted ATP generated from glycolysis (J) or OXPHOS (K) by CTL maintained in amino acid replete media (RPMI) or amino acid deprived (HBSS) with or without VPS34i for 18hr. Data points in bar charts are indicative of biological replicates. Error bars represent the mean +/- SD. P ≤ 0.0001 indicated by ****; P ≤ 0.0005 = ***; P ≤ 0.001 = **; P ≤ 0.05 = *

To test these predictions from the changes in mitochondrial proteins and glucose and lactate transporters, we used a Seahorse extracellular flux analyser. This allowed assessment of the impact of amino acid deprivation in the presence and absence of the VPS34 inhibitor on basal extracellular acidification (ECAR) and oxygen consumption rates (OCR) in CTL. This then allows a gauge of the impact of autophagy induction and repression on rates of ATP generation from glycolysis and oxidative phosphorylation ^50,51^. The data show decreased ECAR from amino acid deprived CTL and a corresponding calculated decreased ATP production from glycolysis (Fig 6J and Supp Fig S6B). When autophagy is repressed by VPS34 inhibition there is substantial ECAR increase and increased production of ATP from glycolysis (Fig 6J and Supp Fig S6B). There is also decreased oxygen consumption by amino acid deprived CTL (Fig S6C) with a major impact on calculated ATP production from oxidative phosphorylation, but these changes were not rescued by VPS34 inhibition (Fig 6K and Supp Fig S6C).

### CTL survival is orchestrated by VPS34 control of autophagy during amino acid starvation

What proteins are fuelled by recycled amino acids from autophagy? These would be proteins whose synthesis was dependent on autophagy, identified as those proteins whose abundance decreased with VPS34 inhibition of autophagy in amino acid deprived CTL (Fig 7A and Supp Table S7). Clustered enrichment analysis highlighted mitochondrial proteins, protein transport, lysosomal and vesicle proteins as well as some involved in DNA binding (Fig 7B, Supp Table S7). The enrichment of mitochondrial proteins is not reflected in an increase of relative mitochondrial mass upon amino acid deprivation (Fig 7C), instead amino acid deprivation drives mitochondrial remodelling of the mitoribosomes, mitoribosome assembly factors (eg ERAL1) and several mitochondrial transporters (Supp Fig S7A, S7B, S7C). Expression of these proteins in response to amino acid starvation is dependent upon VPS34 activity. Other proteins dependent upon autophagy for increased expression include metabolic enzymes GLUL (glutamine synthetase), PIP4K2A and UGCG (UDP-Glucose Ceramide Glucosyltransferase; sphinglolipid metabolism), as well as the arginine transporter SLC7A3 (Fig S7D, S7E), the latter has been shown to be upregulated under conditions of amino acid deprivation in HEK293 and HeLa cells ^52^. The expression of the NFKB inhibitory protein NFKBID and FOXK2 were also dependent on VPS34 activity (Fig S7F). Other noteworthy proteins whose expression was dependent on autophagy in amino acid starved CTL included several proteins associated with cell detoxification ^53–55^ (Fig 7D). The autophagy driven expression of this suite of “protective” proteins raises questions about whether inhibiting autophagy in amino acid deprived CTL will impact cell survival? This was tested experimentally, and a salient result is that inhibition of VPS34 has no impact on cell survival in amino acid replete CTL with low autophagy flux whereas amino acid deprived CTL accumulate high levels of intracellular ROS and cannot survive when VPS34 is inhibited or autophagy blocked by Bafilomycin treatment (Fig 7E, 7F).

**Figure 7:**
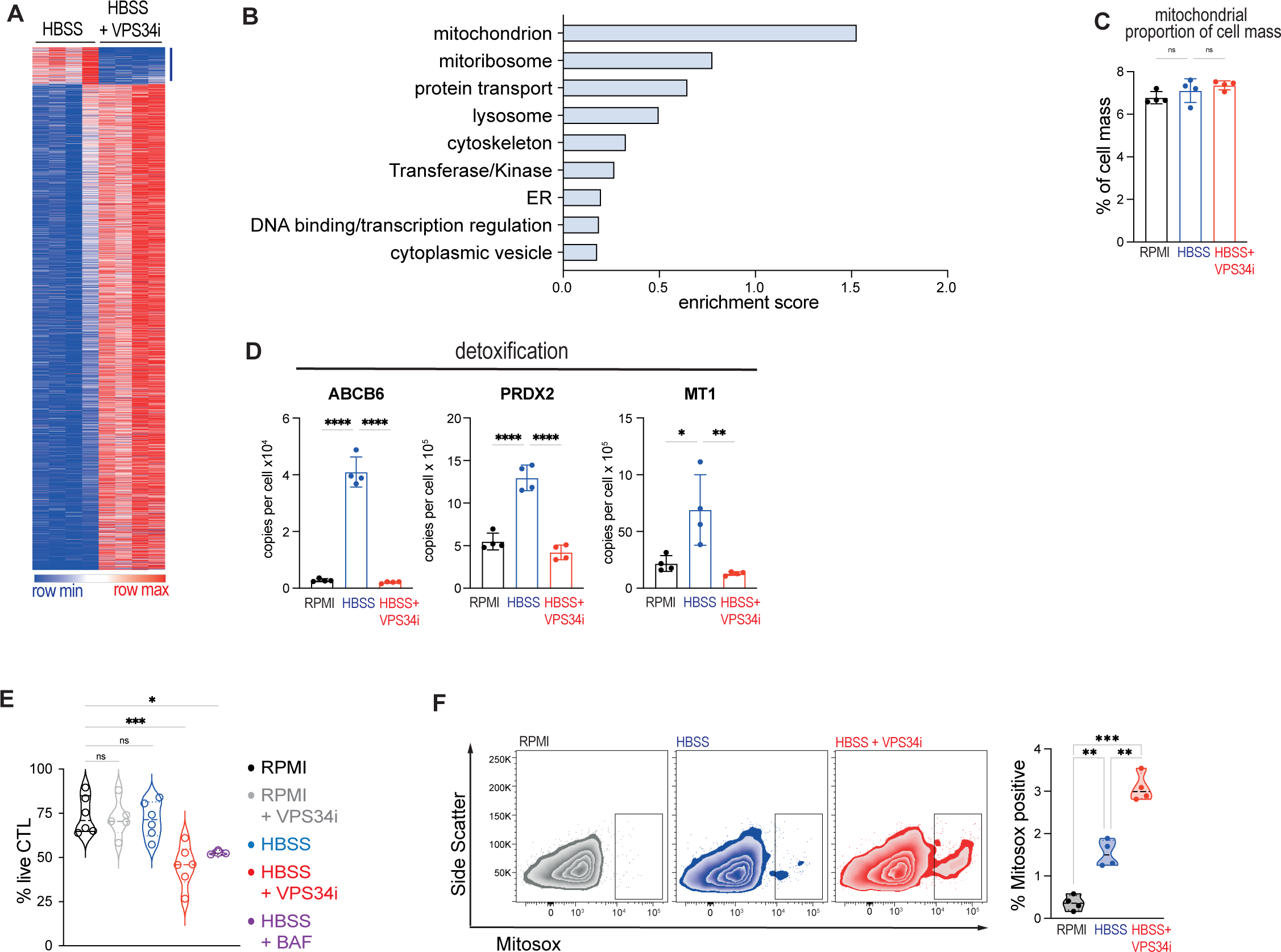
CTL survival program is dependent upon VPS34 during AA starvation. **A** Heatmap of the subset of proteins from CTL which are significantly changed upon amino acid deprivation (HBSS) and with VPS34i. **B** Clustered enrichment analysis on proteins from amino acid deprived (HBSS) CTL that were significantly decreased with VPS34i treatment. The full enrichment table is available in Supplementary data file S7. **C** Protein content of mitochondrial proteins relative to total cell mass. **D** Protein copy numbers per cell of ABCB6, PRXD2 and MT1 from CTL maintained in amino acid replete (RPMI) media or amino acid deprived (HBSS) for 18r with or without VPS34i. **E** Proportion live CTL in replete amino acids (RPMI), or amino acid deprived (HBSS) with or without VPS34i or Bafilomycin A (Baf) for 18hr. **F** Representative flow cytometry data showing Mitosox staining (left) and proportion of Mitosox high cells (right) from CTL in replete amino acids (RPMI) or amino acid deprived (HBSS) with or without VPS34i for 18hr. Data points in bar charts or violin plots are indicative of biological replicates. Error bars represent the mean +/- SD. P ≤ 0.0001 indicated by ****; P ≤ 0.0005 indicated by ***; P ≤ 0.001 indicated by **; P ≤ 0.05 indicated by *

### VPS34 control of autophagy and protein degradation in naïve CD8 T cells

To understand how autophagy shapes naïve CD8 T cells we used mass spectrometry to quantitatively analyse the proteomes of IL-7 maintained naïve T cells cultured in the presence or absence of the VPS34 inhibitor for 5 hours. In these experiments, > 4’900 proteins were quantified and VPS34 inhibition increased the abundance of 327 proteins and decreased expression of 322 proteins, with expression of 35 proteins lost entirely upon VPS34i treatment (Figure 8A and Supp file S8). A clear indication of VPS34 repression of autophagy flux was the increased expression of autophagy cargo adapters p62/SQSTM1, SEC62 and RTN3 and CALCOCO1 in VPS34i treated cells (Fig 8B). Previous studies in have identified the IL-7R as an autophagy cargo in proliferating CD4 T cells such that when autophagy is inhibited surface IL-7 receptor levels accumulate (Zhou et al. 2022). However, VPS34 inhibition did not cause accumulation of IL-7 receptors in naïve CD8 T cells nor did VPS34 inhibition prevent IL-7 induced internalisation of the IL-7 receptor in CD8 T cells (Fig 8C).

**Figure 8:**
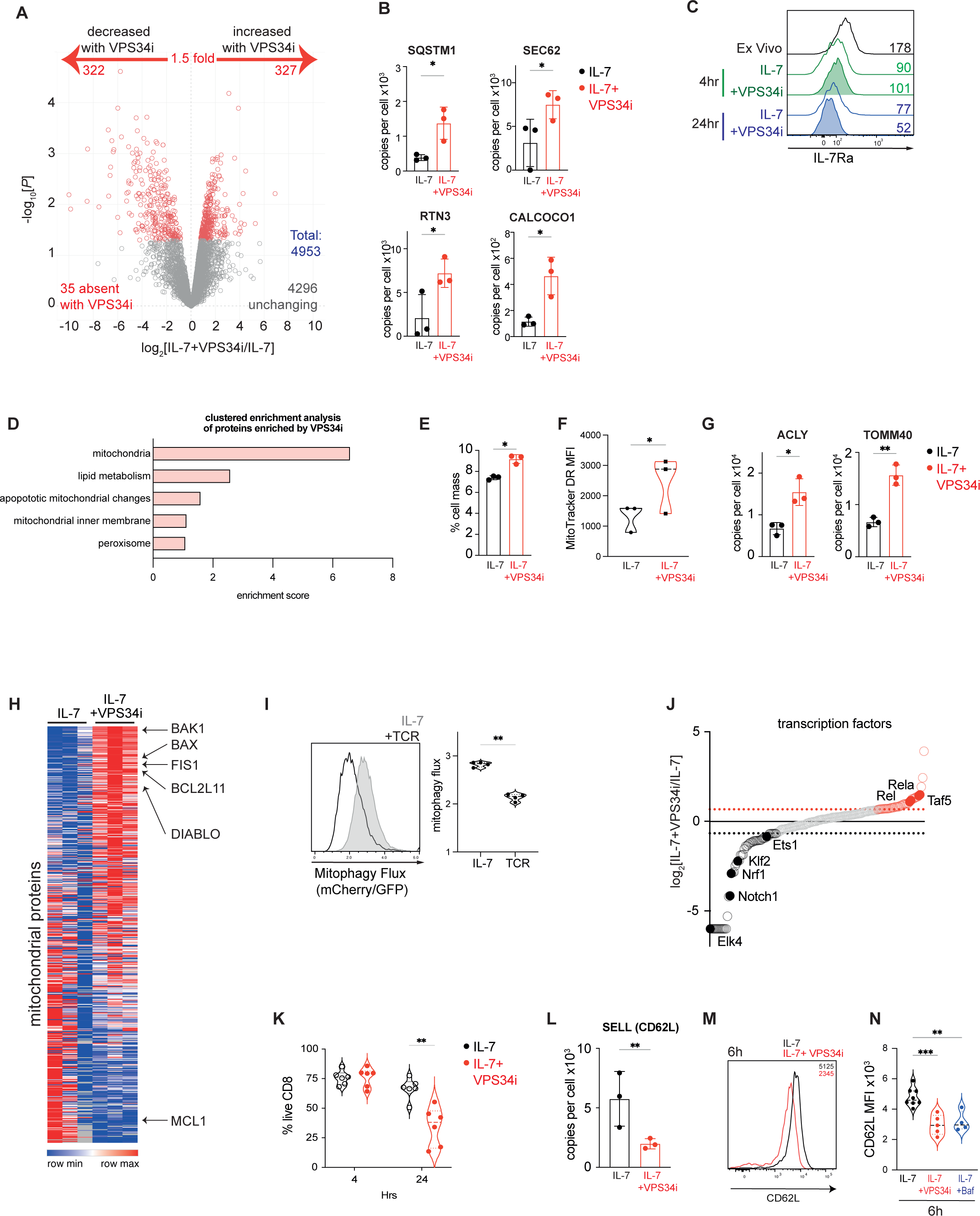
VPS34 control of autophagy and protein degradation in naïve CD8 T cells. **A** Volcano plot showing the ratio changes of proteins in IL-7 maintained CD8 T cells treated with VPS34i for 5 hrs. *P* values below 0.05 and fold change above 1.5 are considered significant and marked in red. The full list of proteins detected is available in Supplementary Table S8. **B** Protein copy numbers per cell of SQSTM1, SEC62, RTN3 and CALCOCO1. **C** Representative flow cytometry staining of IL-7Ra expression on CD8 T cells either ex vivo, or maintained in IL-7 with or without VPS34i for 4hr or 24hr. MFI values are presented in the plot. **D** Clustered enrichment analysis of proteins increased in naïve CD8 T cells treated with VPS34i for 5 hrs. The full enrichment table is available in Supplementary data file S8. **E** Protein content of mitochondrial proteins relative to total cell mass. **F** Flow cytometry mean fluorescence intensity (MFI) values of MitoTracker DeepRed staining of CD8 T cells maintained in IL-7 with or without VPS34i for 5hr. **G** Protein copy numbers per cell of ACLY and TOMM40 from IL-7 maintained CD8 T cells +/- VPS34i for 5 hrs. **H** Heatmap of mitochondrial proteins (GO 0005739, MitoCarta3.0). **I** Representative flow cytometry histogram showing Mitophagy flux (mCherry/GFP) profiles of mitophagy reporter CD8 cells maintained in IL-7 or activated through the TCR (aCD3/CD28) for 6hr (left) and corresponding mean Mitophagy flux (mCherry/GFP) values (right). **J** Ranked expression of transcription factors (GO 00037000) detected by proteomics. 1.5 fold decreased with VPS34i are shown in black, and 1.5 fold increased with VPS34i are shown in red. The full enrichment list is available in Supplementary data file S8. **K** Proportion live CD8 T cells maintained in IL-7 with or without VPS34i for 4hr or 24hr. **L** Protein copy numbers per cell of SELL (CD62L) from IL-7 maintained CD8 T cells +/- VPS34i for 5 hrs. **M** Representative flow cytometry staining of CD62L surface expression on CD8 T cells maintained in IL-7 with or without VPS34i for 6hr, corresponding MFI values are presented in the plot. **N** Flow cytometry MFI values of CD62L surface staining from CD8 T cells maintained in IL-7 +/- VPS34i or Bafilomycin A (Baf) for 6hr. Data points in bar charts or violin plots are indicative of biological replicates. Error bars represent the mean +/- SD. P ≤ 0.0001 indicated by ****; P ≤ 0.0005 = ***; P ≤ 0.001 = **; P ≤ 0.05 = *.

Clustered enrichment analysis indicated that the dominant effect of VPS34 inhibition in naïve T cells was to allow accumulation of mitochondrial proteins (Fig 8D and Supp table S8). Both mitochondrial mass as a proportion of total cellular mass (Fig 8E) and direct mitochondrial labelling with Mitotracker DeepRed (Fig 8F) show increased mitochondrial protein levels after VPS34 inhibition. Structural proteins such as membrane transporters, mitochondrial ribosomal subunits and mitochondrial enzymes (eg ACLY, TOMM40) all accumulate in response to VPS34 inhibition (Fig 8G and Supp table S8, Supp file S8). This is consistent with previous studies showing that VPS34 null T cells increase mitochondrial mass (Willinger and Flavell 2012). However, the insight from the present study is the striking selectivity of the mitochondrial proteome remodelling that occurs upon VPS34 inhibition (Fig 8H). One notable observation was that VPS34 inhibitor treated naïve CD8 T cells accumulate the pro-apoptotic mitochondrial pore proteins BAK1, BIM (BCL2L11), DIABLO and BAX and the mitochondrial fission-related protein FIS1 (Fig 8H). These data argue that the accumulation of mitochondrial proteins after autophagy suppression reflects accumulation of abnormal mitochondria. In this context, the suppression of autophagy flux in naïve T cells following antigen receptor engagement correlated with accumulation of BAK1 and BAX (Supp Fig S8A). These data suggest that a dominant form of autophagy in naïve T cells is mitophagy. Accordingly, TCR-induced suppression of autophagy flux in naïve CD8 T cells is actually TCR-suppression of mitophagy. To test this hypothesis, we made use of the *mito*-QC transgenic mitophagy reporter ^21,22,56^. Mitochondria from this mouse are decorated with a tandem mCherry-GFP tag localised to the outer mitochondrial membrane, and as with the autophagy reporter, mitophagy flux can be monitored by calculating GFP: mCherry fluorescence ratios. The data in figure 8I confirm that naïve CD8 T cells have high levels of mitophagy flux, which is decreased upon TCR activation.

We also considered what proteins might be fuelled by amino acids recycled by autophagy in naïve T cells. Here, candidates are the ∼350 proteins whose abundance was decreased or absent with VPS34 inhibition (Fig 8A, supplemental table). Notably, VPS34 inhibition caused decreased expression of KLF2, ETS family and Nrf1 transcription factors (Fig 8J, Supp Fig 8B) and the anti-apoptotic family member MCL-1 (Fig 8H). The loss of MCL-1 predicts that sustained inhibition of autophagy in naïve CD8 T cells would impact cell survival because MCL-1 has been shown to mediate pro-survival responses of IL7 in naive T cells. When this hypothesis was tested experimentally it was observed that VPS34 inhibition of IL7 maintained naïve CD8 T cells for 24 hours results in their increased death (Fig 8K). One other salient observation was that VPS34 inhibition decreased the abundance of CD62L (SELL, L-selectin, CD62L), a critical adhesion receptor that controls naive T cell trafficking into secondary lymphoid tissues (Fig 8L) ^57–59^. This observation was tested orthogonally using flow cytometry which also showed decreased expression of CD62L in VPS34 inhibitor treated naïve CD8 cells (Fig 8M, 8N). It has been shown that CD62L has a short half-life in naive T cells and would need to be constantly synthesized to maintain membrane expression (Wolf et al. 2020); the present data indicate that VPS34 mediated autophagy in naïve T cells produces amino acids that fuel CD62L synthesis.

## Discussion

The present study provides new insights about autophagy regulation during T cell immune responses that revises many previous ideas about autophagy timing in relation to T cell immune activation. Fundamental discoveries are that autophagy, particularly mitophagy, is high in naïve T cells and that immune activation represses mitophagy flux and represses protein degradation via autophagy. If CD8 T cells differentiate to effectors in response to inflammatory cytokines such as IL-2, IL-12, and IL-18, autophagic flux and protein degradation by autophagy is repressed. However, amino acid or cytokine deprived effector T cells rapidly revert to high levels of autophagic protein degradation. As well, antigen activated cells differentiated to memory CD8 T cells in response the cytokine IL-15 retain high levels of autophagy flux. These present data are consistent with observations of decreased autophagy in T cells proliferating *in vivo* in response to viral infection ^7^. Moreover, high levels of autophagy in naive and memory cells reported herein fits with gene deletion studies that showed that the loss of autophagy regulators during T cell development causes loss of these populations *in vivo* ^1–9^.

Many previous studies have concluded that immune activation of T cells increases autophagy ^1,2,13–15^. These conclusions are often based on quantification of autophagosomes or levels of lipidated LC3b ^16–18^. Our conclusions about autophagy flux are based on the use of a sensitive autophagy flux reporter ^21^ as well as mass spectrometry-based analysis of autophagy dependent protein degradation in different T cell populations. Our data reveal that levels of multiple components of the autophagic machine are indeed highly increased in immune activated cells. For example, LC3b is only detected in effector but not naïve cells. However, the new insight is that increases in expression of autophagy machinery in activated T cells occurs because autophagy flux is repressed by antigen receptor triggering and inflammatory cytokines such that the autophagy machinery is not degraded and accumulates.

To fully understand why this pattern of autophagy regulation has evolved it is of value to understand the proteins and organelles that are selected for autophagic degradation in naïve or effector CD8 T cells. Here, we used mass spectrometry to census the protein targets for autophagy. A salient fact to emerge from these experiments is that there are fundamental differences in autophagy cargoes in different CD8 populations. Hence, we show that the dominant cargoes for degradation by autophagy in naïve cells are mitochondrial proteins and we show that naïve T cells use mitophagy to prune mitochondria to maintain mitochondrial integrity. In contrast, in cytotoxic T cells when autophagy is induced by nutrient deprivation, the dominant proteins degraded by autophagy are not mitochondria: indeed the data indicate that the induction of autophagy in response to amino acid deprivation results in remodelling of mitochondria using amino acids produced by autophagy. The dominant autophagy substrates in effector T cells are the contents of cytotoxic granules such as perforin and granzymes. Other proteins degraded by autophagy in effector CD8 T cells are key metabolic proteins such as glucose, lactate and amino acid transporters and glycolytic enzymes. Effector T cells thus use autophagy to downregulate energy consuming processes to switch to energy conserving pathways to promote survival. Herein we show that cytokine deprived T cells undergoing autophagy downregulate their cytolytic capacity, but this can be prevented by inhibition of autophagy. Collectively, these observations afford an explanation for why inactivation of T cell autophagy has been shown to enhance tumour rejection and why T cells deficient in autophagy increase glucose transport and metabolism ^11,12^. The current data also provide a molecular explanation for why autophagy is important for naïve T cell homeostasis ^5^. Hence, in the absence of autophagy, naïve T cells accumulate damaged mitochondria which would promote apoptosis. As well, when autophagy is inhibited, naïve T cells cannot sustain expression of CD62L a crucial adhesion molecule which ensures naïve T cell trafficking to secondary lymphoid tissues ^57,58^. In naive T cells, CD62L is constantly synthesized, trafficked to the membrane but then continually cleaved and shed with half-life of approximately 1 hour ^60–62^. The dependence of CD62L expression on autophagy indicates that CD62L synthesis in naive cells uses amino acids from proteins degraded by autophagy. Decreased CD62L expression in T cells lacking autophagy would prevent T cell trafficking to secondary lymphoid tissues.

One other key finding is that autophagy flux in T cells is very dynamic can change rapidly in response to immune stimulation and or nutrient deprivation. In this context, we show that autophagy flux in T cells is controlled by amino-acid sensing. Nutrient control of autophagy is evolutionarily conserved for yeast to mammals but what is perhaps unique to immune cells is that the ability of T cell to ‘sense’ amino acids is regulated by the restriction of amino acid transporter expression to immune activated T cells. The default state of high autophagy/mitophagy in naïve or memory T cells reflects that these have low levels of amino acid transporters and hence cannot amino-acid sense their environment. We show that autophagy repression occurs when T cells respond to immune activation by antigen receptors and proinflammatory cytokines to upregulate and sustain amino acid transporter expression. High levels of amino acid transporters on activated effector T cells thus allows them to repress autophagy when they are in an amino acid replete environment. Importantly, the differential ability of cytokines to repress autophagy reflects the differential ability of some cytokines to control amino acid transporter expression. For example, IL-2 and 12 which drive effector differentiation repress autophagy because they induce high levels of amino acid transport. In contrast, IL-15 which supports memory T cells cannot repress autophagy because it only supports very low levels of amino acid transporter expression. The importance of the regulated expression of amino acid transporters for the coordination of protein synthesis during T cell immune responses is clear. The new insight is that the immune regulation of amino acid transport capacity is a key autophagy checkpoint that affords a molecular mechanism for antigen receptors and inflammatory cytokines to repress autophagy flux, one of the major protein degradation pathways in T cells. The repression of this energetically demanding protein degradation process would allow a cell to divert energy to support de novo synthesis of proteins that support T cell cytokine production, clonal expansion and differentiation rather than using ATP to replenish degraded proteins.

## Materials and Methods

### Mice and cells

OT-1 mice (Charles river UK, stock number 003831)^68^; autophagy reporter mice ^21^; mitophagy reporter/mito-QC transgenic mice ^21,22,56^ and C57BL/6 (wild-type, WT) mice were bred and maintained in the WTB/RUTG, Univeristy of Dundee in compliance with UK Home Office Animals (Scientific Procedures) Act 1986 guidelines.

T cells were cultured in complete culture medium - RPMI 1640 containing glutamine (Gibco), supplemented with 10 % FBS (Gibco), 1 % penicillin/streptomycin (Gibco) and 50 µM β-mercaptoethanol (Sigma) at 37°C and 5% CO_2_. Single cell suspensions were generated by disaggregation of lymph nodes or spleens. Splenic red blood cells were lysed in ACK buffer (150 mM NH_4_Cl, 10 mM KHCO_3_, 110 µM Na_2_EDTA, pH 7.8). Cell suspensions were washed in complete culture medium (RPMI 1640 containing glutamine (Gibco), supplemented with 10 % FBS (Gibco), 1 % penicillin/streptomycin (Gibco) and 50 µM β-mercaptoethanol (Sigma)) prior to activation. To maintain naïve T cells, splenic and lymph node single cell suspensions were cultured with Interleukin 7 (IL-7) (5 ng/ml, PeproTech) in complete RPMI 1640 medium. VPS34-IN1 (VPS34i) (1nM), Halofuginone (100nM), Bafilomycin A1 (200nM), ruxolitinib (1uM) or rapamycin (20nM) were used where indicated.

The EasySep Mouse CD8+ T Cell Isolation Kit (STEMCell technologies) was used to purify naïve OT-1 CD8+ T cells for proteomic analysis.

### Polyclonal T cell activation

To generate cytotoxic T lymphocytes (CTL), cells were stimulated with 1ug/ml of anti-CD3 (2C11, BioLegend) and 2ug/ml anti-CD28 (37.51, ebiosciences) in complete media for 48hrs. Cells were then washed out of activation conditions, expanded and split daily into fresh complete RPMI 1640 medium with IL-2 (201ng/ml, Proleukin, Novartis) at a density of 3×10^5^ cells/ml for a further 3–5 days.

### OT-I TCR transgenic T cell activation

For generation of OT-1 CTL, lymph nodes single cell suspensions were isolated from OT-1 TCR transgenic mouse and stimulated for 36 hours with SIINFEKL peptide (final concentration of 10ng/ml). Cells were then washed out of activation conditions expanded and split daily into fresh complete RPMI 1640 medium with IL-2 (201ng/ml, Proleukin, Novartis) at a density of 3×10^5^ cells/ml for a further 3–5 days.

For experiments with IL-12 and IL-18, activated OT-1 T cells were initially expanded in IL-2 (20ng/ml) and IL-12 (20ng/ml; Peprotech) before being switched into IL-2 alone, IL-12 + IL-18 (20ng/ml; R&D Systems) or no cytokine for 24h. For experiments with IL-15, activated OT-1 T cells were expanded in IL-15 (20ng/ml; Peprotech).

### Nutrient deprivation studies

Prior to nutrient deprivation, cells were washed twice in PBS. Cells were then resuspended at 3×10^5^ cells/ml in either complete RPMI or nutrient deprivation medium listed below. All nutrient deprivation media were supplemented with 10 % Dialysed FBS (dFCS) (Gibco), 1% penicillin/streptomycin (Gibco) and 50 µM β-mercaptoethanol (Sigma).

- For total amino acid starvation cells were cultured in Hanks’ Balanced Salt solution (HBSS, Gibco).
- For glucose starvation cells were cultured in RPMI 1640 media with no glucose (Gibco).
- For glutamine starvation cells were cultured in RPMI 1640 media lacking glutamine (Gibco).
- For arginine or methionine starvation cells were cultured in RPMI 1640 media lacking arginine, lysine and methionine (DC Biosciences). Supplementation of individual amino acids (Sigma) to the levels of standard RPMI 1640 formula was carried out to generate culture medium lacking a single amino acid, methionine or arginine.

### Flow cytometry sample acquisition and analysis

For surface staining, antibody clones used were: CD8a (53-6.7), CD44 (IM7), IL-7Ra (CLONE), CD62L (MEL-14), Fc Block (BD Biosciences). Antibodies conjugated to APC, AlexaFluor 647, PeCy7, PerCPCy5.5, APC-efluor780, Brilliant Violet 421 and 605 were obtained from either eBioscience, Biolegend or BD Biosciences.

For intracellular staining, cells were fixed with 1% formaldehyde (v/v) prior to permeabilization with 90% (v/v) ice cold methanol for 30 mins. Cells were washed and incubated with antibody against phospho S6 Ser235/236 (cat no. 2211) or phospho ACC S79 (clone D7D11 cat no. 11818) prior to incubation with anti-rabbit Alexa 647 secondary (cat no. 4414; all Cell Signaling technology). Cells were washed and resupended in 0.5% FBS/PBS (v/v) for flow cytometric acquisition.

Mitotracker Deep Red (cat no. M22426) and MitoSOX (cat no. M36008) staining was performed according to manufacturer’s guidelines (Invitrogen, Thermo Scientific).

Kynurenine uptake to monitor System L transporter activity was done according to established protocols ^69^. Briefly, surface stained cells were incubated in pre-warmed HBSS (GIBCO) with 200uM kynurenine (Sigma) for 4 min at 37C prior to fixation with 1% formaldehyde (v/v). Where indicated kynurenine uptake was performed with 10mM BCH (Sigma), a competitive inhibitor for System L uptake. Cells were washed post-fixation and resuspended in 0.5% FBS/PBS for flow cytometric acquisition.

Flow cytometry was performed on FACSVerse, or BD LSRFortessa flow cytometers (BD Biosciences). During acquisition, lymphocytes were identified by gating on forward scatter area (FSC-A) and side scatter area (SSC-A). Doublets were excluded using FSC-A vs forward scatter width (FSC-W). Specific experimental gating strategies are referred to in figure legends. FlowJo analysis software V10.9 and V10.10 (Becton Dickinson) was used to analyse all flow cytometry experiments.

### Flow cytometric analysis of autophagy reporter T cells

When cells are not undergoing autophagy the mCherry: GFP:LC3b displays a diffuse cytosolic staining pattern, with both GFP and mCherry fluorescence. Upon autophagosome-lysosome fusion, GFP fluorescence becomes quenched by the low pH whilst mCherry is not affected. Autophagy flux can thus be measured by calculating the GFP:mCherry fluorescence ratio as a derived parameter in FlowJo. GFP or mCherry expressing Jurkat cells were used as compensation controls for flow cytometry.

### Listeria monocytogenes infection model

Mice were intravenously infected with 1-2 x 10^6^ colony forming units (CFU) of attenuated ActA-deficient *L. monocytogenes* ^70^. On days 1, 3 and 7 post-infection spleens were harvested from 3 infected mice and 1 uninfected control mouse and analysed by flow cytometry.

### CTL killing assay

OT-1 CTL were maintained in IL-2 (20ng/ml) or deprived of IL-2 with or without VPS34i (1nM) for 24h prior to co-culture with E.G7-OVA (ATCC, CRL-2113) target cells in a 5:1 Target:T ratio. E.G7-OVA target cells were stained using CellTracker-DeepRed (ThermoFisher), according to manufacturer’s instructions, 24h prior to the killing assay. The killing capacity of the CTL is represented by the % of target cells undergoing apoptosis after 4hr: the co-culture was stained with CellEvent-Caspase3/7-Green (ThermoFisher) for 30mins, then fixed in IC Fixation Buffer (Invitrogen), 30mins at RT, then washed twice in PBS and analysed with an iQue Screener (Sartorius). CellTracker-stained cells with high intensity CellEvent-Caspase3/7-Green were determined to be apoptotic target cells.

### Metabolic assay

The oxygen consumption rate (OCR) and extracellular acidification rate (ECAR) of CTL, either maintained with amino acids (complete RPMI) or 18hr starved of amino acids (HBSS) +/- VPS34i, were measured using a Seahorse XF24 analyser and following established protocols ^35,50,71^. Briefly, 2e5 CTL were plated in poly-D-lysine coated XF24 plates and gently centrifuged in appropriate XF medium. Data was normalised to cell number, and basal and oligomycin treated measurements were used to determine ATP production via OXPHOS or glycolysis based as described in Mookerjee et al ^51^.

### Proteomics sample preparation

Cell pellets were prepared for mass spectrometry analysis as described previously ^72^. In brief, cells were lysed in 400 μl lysis buffer (5% SDS, 10 mM tris(2-carboxyethyl)phosphine, 50 mM Triethylammonium bicarbonate) and incubated for 5 minutes at room temperature shaking at 1000rpm. Samples were then incubated at 95°C for 5 minutes at 500rpm, and then shaken again at room temperature as described above. Samples were sonicated for 15 cycles of 30 s on/ 30 s off with a BioRuptor (Diagenode). To remove trace DNA contamination 1 μl benzonase was added to each sample followed by incubation at 37°C for 15 minutes. Protein concentration was determined using the EZQ quantification kit (Thermo Fisher Scientific). Samples were then alkylated with the addition of iodoacetamide to a final concentration of 20 mM and incubated for 1 hour in the dark at 22 °C. Protein lysates were processed using S-Trap mini columns (Protifi) following the manufacturers instructions. Eluted peptides were dried overnight before being resuspended in 50 µl 1% formic acid and peptide concentration measured using the CBQCA assay following manufacturers protocol (Thermo Fisher Scientific).

### Mass spectrometry

Peptides generated from IL7 maintained naïve cells and CTL were analysed by data independent acquisition (DIA) on a Q Exactive HF-X (Thermo Scientific) mass spectrometer coupled with a Dionex Ultimate 3000 RS (Thermo Scientific). 1.5 µg of peptide from each sample was analysed as described previously ^73^. LC buffers were the following: buffer A (0.1% formic acid in Milli-Q water (v/v)) and buffer B (80% acetonitrile and 0.1% formic acid in Milli-Q water (v/v)). 1.5 μg aliquot of each sample was loaded at 15 μL/min onto a trap column (100 μm × 2 cm, PepMap nanoViper C18 column, 5 μm, 100 Å, Thermo Scientific) equilibrated in 0.1% trifluoroacetic acid (TFA). The trap column was washed for 3 min at the same flow rate with 0.1% TFA then switched in-line with a Thermo Scientific, resolving C18 column (75 μm × 50 cm, PepMap RSLC C18 column, 2 μm, 100 Å). The peptides were eluted from the column at a constant flow rate of 300 nl/min with a linear gradient from 3% buffer B to 6% buffer B in 5 min, then from 6% buffer B to 35% buffer B in 115 min, and finally to 80% buffer B within 7 min. The column was then washed with 80% buffer B for 4 min and re-equilibrated in 3% buffer B for 15 min. Two blanks were run between each sample to reduce carry-over. The column was kept at a constant temperature of 50°C at all times.

The data was acquired using an easy spray source operated in positive mode with spray voltage at 1.9 kV, the capillary temperature at 250°C and the funnel RF at 60°C. The MS was operated in data-independent acquisition (DIA). A scan cycle comprised a full MS scan (m/z range from 350–1650, with a maximum ion injection time of 20 ms, a resolution of 120 000 and automatic gain control (AGC) value of 5 × 106). MS survey scan was followed by MS/MS DIA scan events using the following parameters: default charge state of 3, resolution 30.000, maximum ion injection time 55 ms, AGC 3 × 106, stepped normalized collision energy 25.5, 27 and 30, fixed first mass 200 m/z. Data for both MS and MS/MS scans were acquired in profile mode. Mass accuracy was checked before the start of samples analysis.

### Mass spectrometry data analysis

Raw DIA mass spec data files were searched using Spectronaut version 16.0.220606.53000 (Biognosys). Data was analysed using a hybrid library approach. For IL7 maintained cells the library was assembled using a deep proteome of naïve CD8 T cells along with the experimental DIA samples. For CTL, the library was assembled using a deep proteome of CTL along with experimental DIA samples. Naïve and CTL deep proteomes were generated by fractionating peptides from these samples using high pH reverse phase fractionation and analysing by data dependent acquisition. 1 µg of peptide from each fraction was analysed using a LTQ Orbitrap Velos (Thermo Fisher Scientific) as described in detail previously ^74^. Raw mass spec library data files were searched using the Pulsar tool within Spectronaut using the following settings and as described previously ^72^: 0.01 FDR at the protein and peptide level with digest rule set to ‘TrypsinP’. A maximum of two missed cleavages and minimum peptide length of 7 amino acids was selected. Carbamidomethyl of cysteine was selected as a fixed modification while protein n-terminal acetylation and methionine oxidation were selected as variable modifications. The data was searched against a mouse database from the Uniprot release June 2020. The database was generated using all manually annotated mouse SwissProt entries, combined with mouse TrEMBL entries with protein level evidence available and a manually annotated homologue within the human SwissProt database. Experimental DIA samples were then searched against both the fractionated DDA library and the experimental DIA library using the following identification settings: protein and precursor q-value set to 0.01 with an ‘Inverse’ decoy method and ‘Dynamic’ decoy limit strategy. The following quantification settings were used: ‘Quant 2.0’, the MS-Level quantity was set to ‘MS2’, imputation was disabled, major group Top N and minor group Top N were set as ‘False’ and cross run normalisation was set as ‘False’.

Protein copy numbers per cell were estimated using the “proteomic ruler” ^75^. P-values and fold changes for volcano plots were calculated using RStudio (v 2023.06.1+524) with the Bioconductor package “limma” (v3.54.2) ^76^. Heatmaps were generated using the Morpheus data visualization tool hosted by the Broad Institute (Morpheus, https://software.broadinstitute.org/morpheus).

Mass spectrometry raw data and search files will be publicly available via ProteomeXchange and proteomics data will be freely available on the Immunological Proteome resource (ImmPRes, immpres.co.uk) upon publication.

### Statistical analysis

Data are presented as mean values ± standard deviation unless otherwise stated. Statistical analyses were performed using Prism 10, GraphPad Software or Microsoft Excel. An Unpaired t-test was used to compare different media or cytokines. A Paired t-test was used to compare treatment with or without inhibitor. Multiple comparisons in one-way ANOVA analyses were corrected using the Tukey method. The Shapiro-Wilk test for normality was used to determine suitable tests for parametric or non-parametric populations. Alternative tests used are stated in the respective figure legends.

Gene Ontology (Go) term analysis and Kyoto Encyclopaedia of Genes and Genomes (KEGG) pathway analysis was completed by using the online software tool DAVID (v2023q4) ^77,78^.

## Declarations

All procedures involving animals were approved by the University of Dundee Ethical Review Committee and under the authorisation of the UK Home Office Animals (Scientific Procedures) Act 1986.

## Supporting information

Supplemental Datafile S5

Supplemental Datafile S6

Supplemental Datafile S7

Supplemental Datafile 8

