## Supplementary figures and figure legends for "Autophagy repression by antigen and cytokines shapes mitochondrial, migration and effector machinery in CD8 T cells"

**Supplemental Figure S1**

S1A

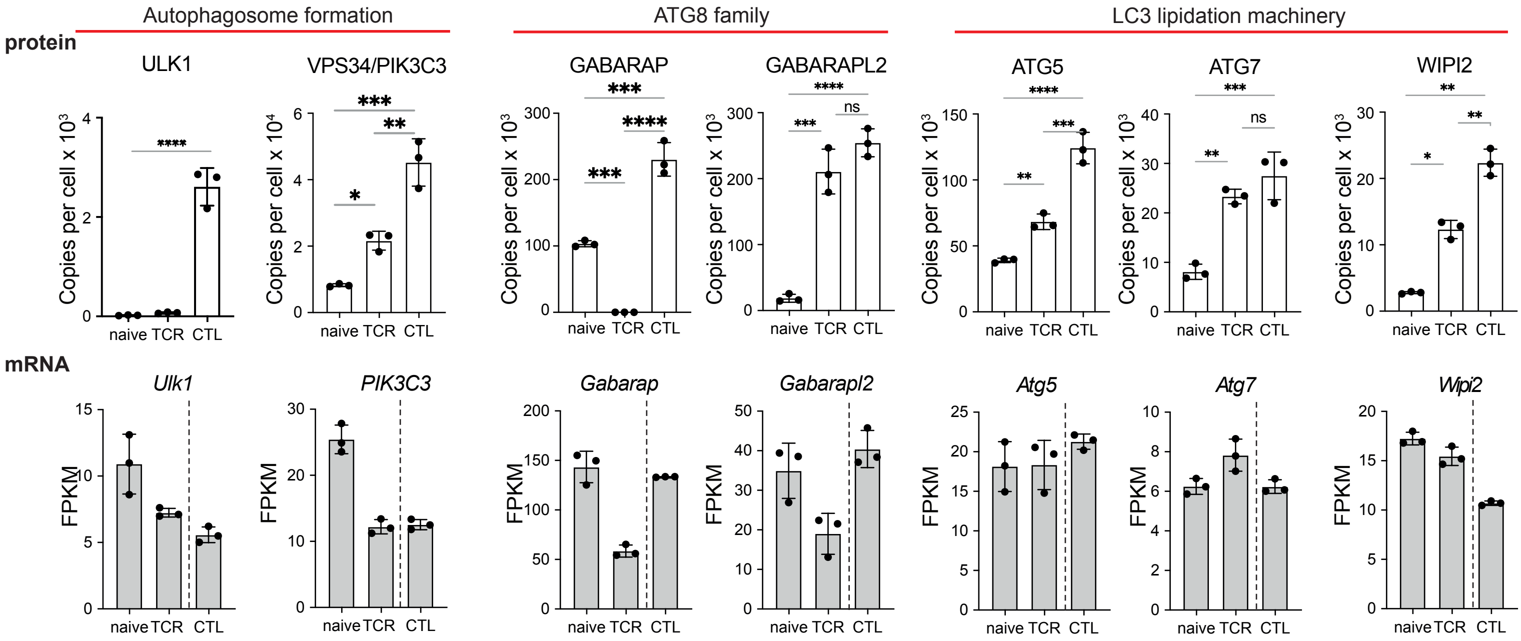


S1B

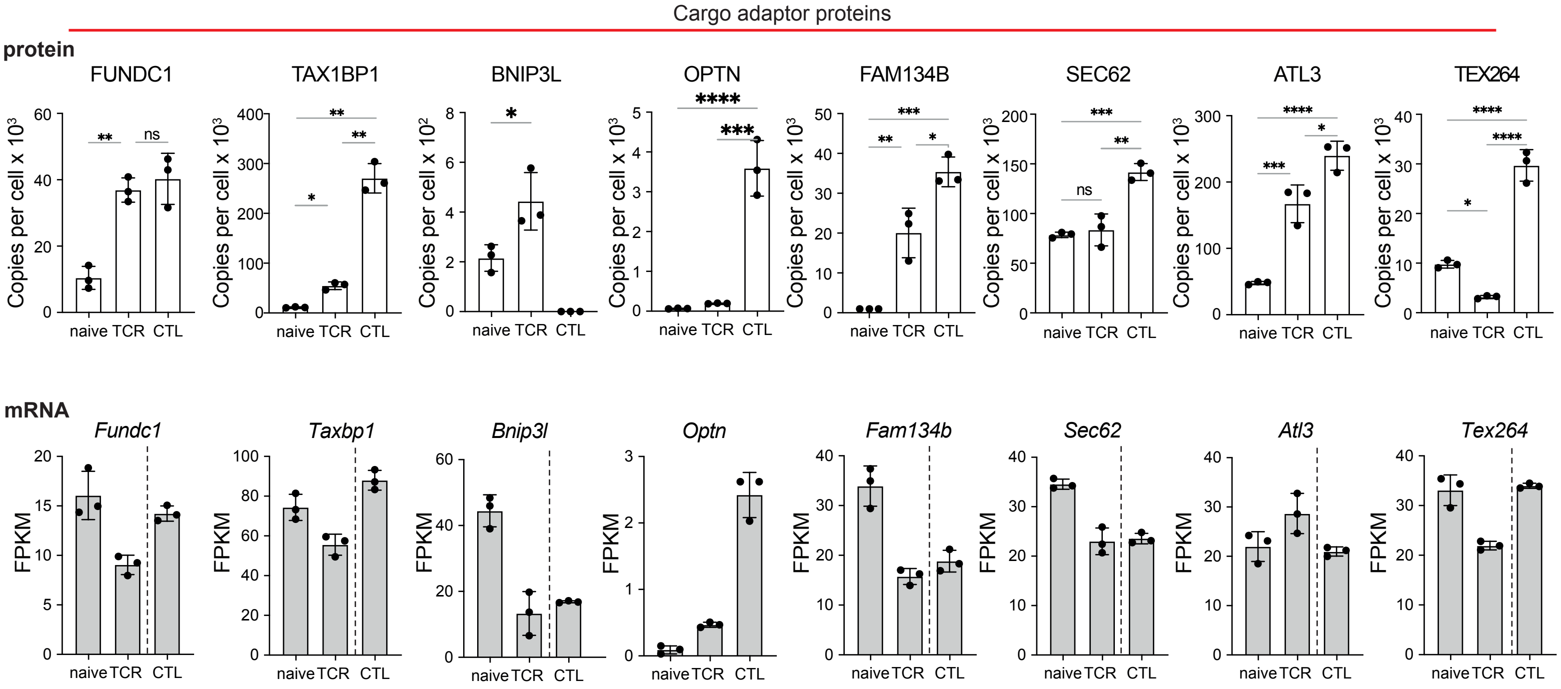


S1C

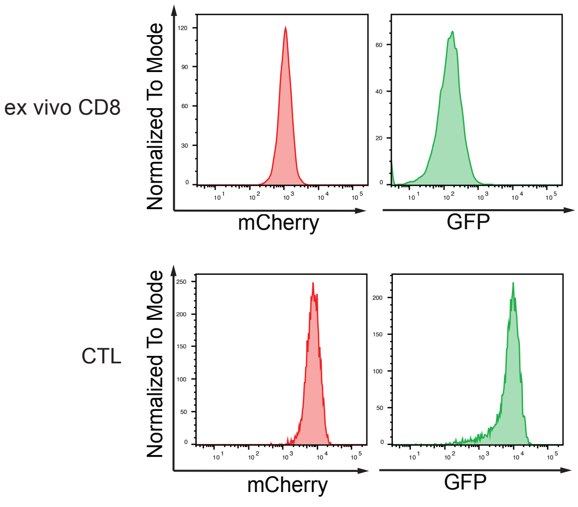

**Supplemental Figure S2**

## S2A

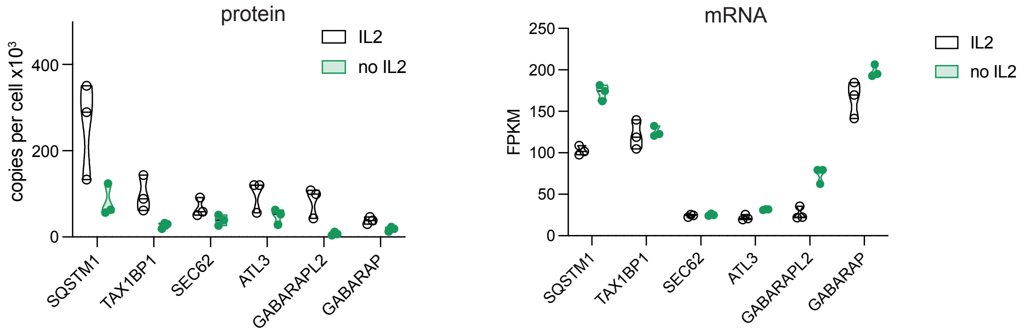


## S2B

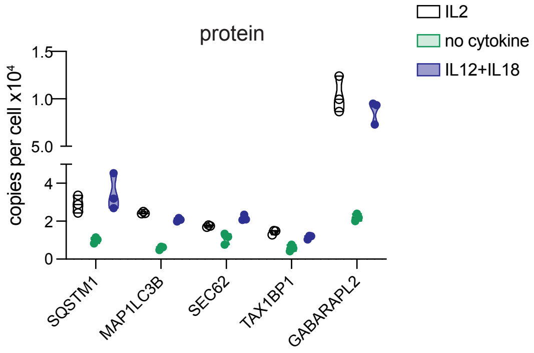


## S2C

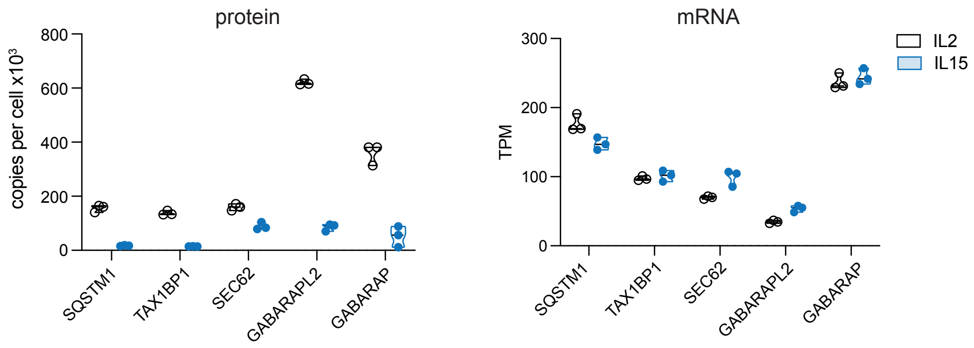

Supplemental Figure S3

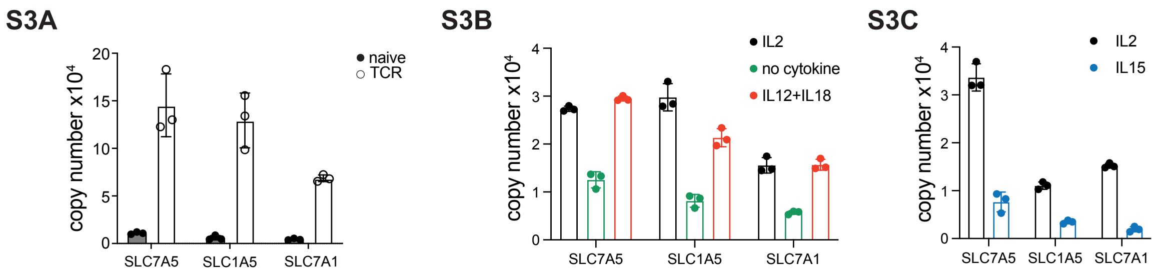

**Supplemental Figure S4**

S4A

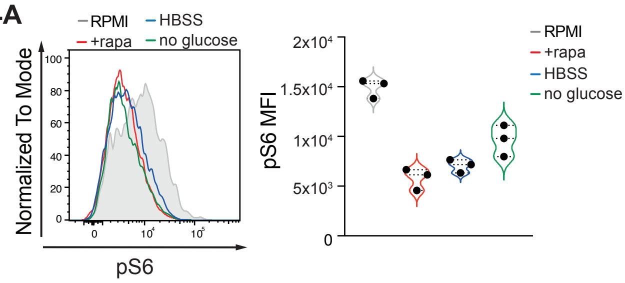


S4B

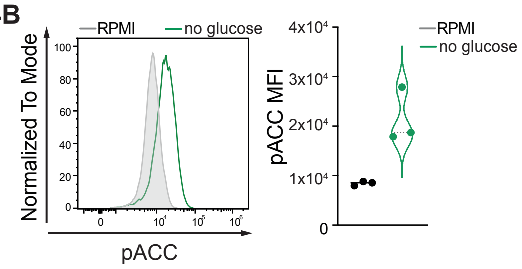

Supplemental Figure S5

S5A

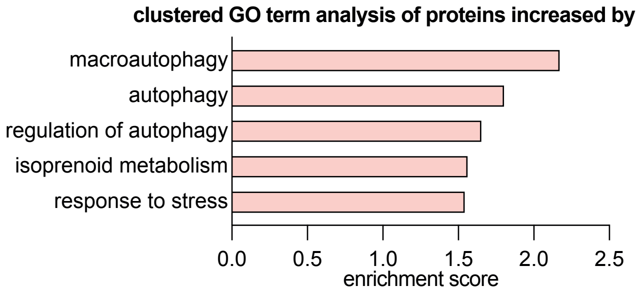


S5B

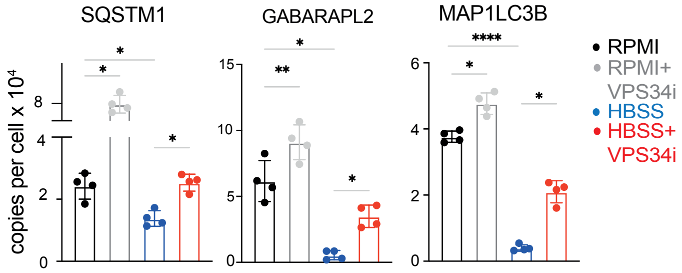


S5C

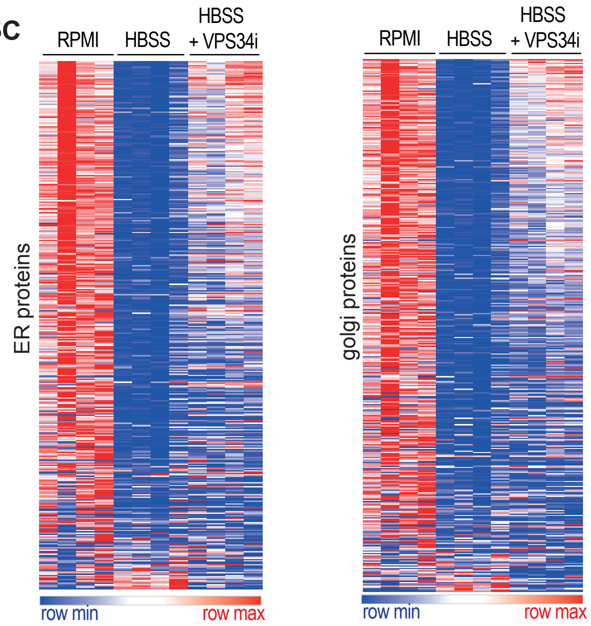


S5D

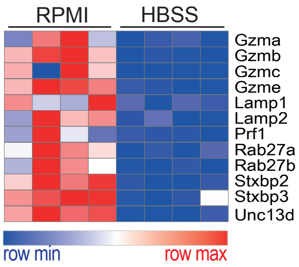


S5E

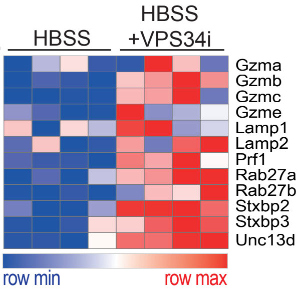

**Supplemental Figure S6**

S6A

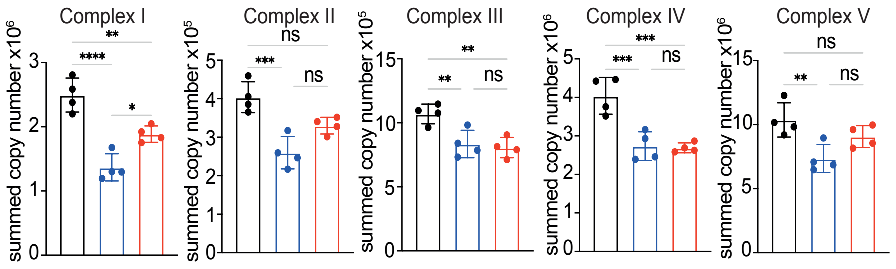


S6B

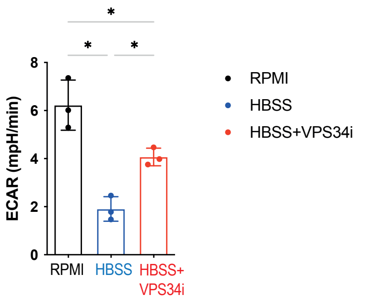


S6C

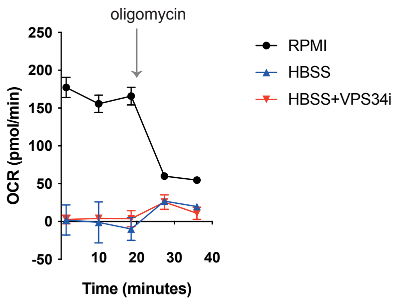

**Supplemental Figure S7**

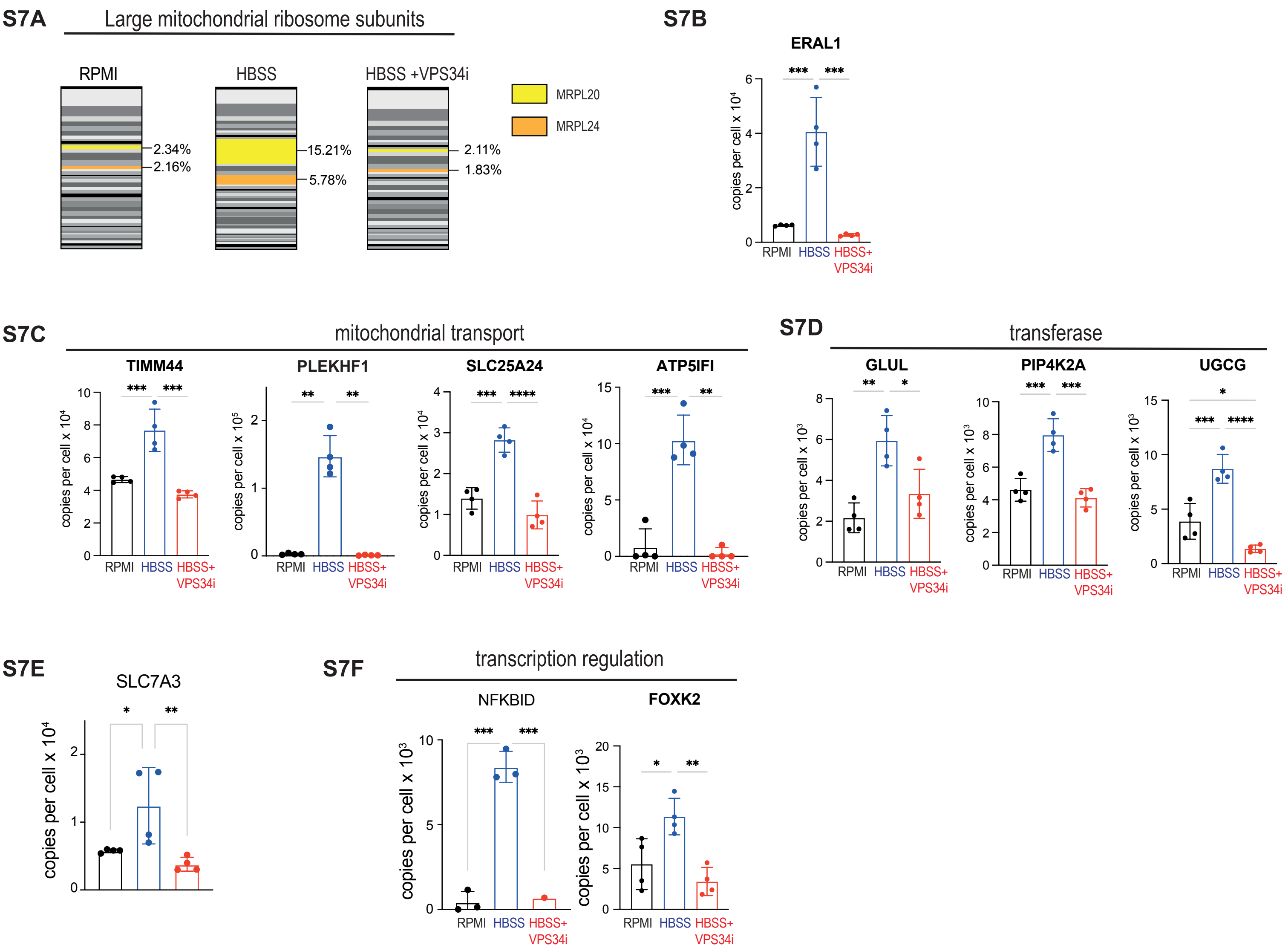

**Supplemental Figure S8**

**S8A**

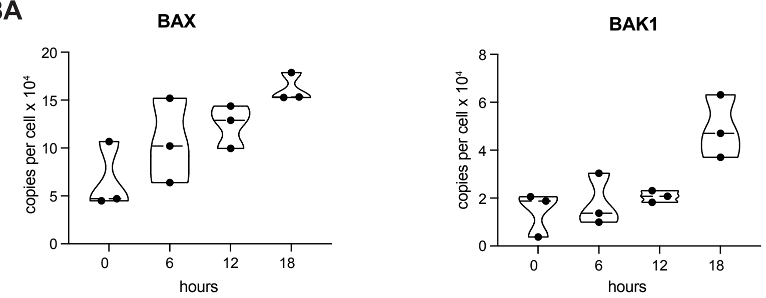


**S8B**

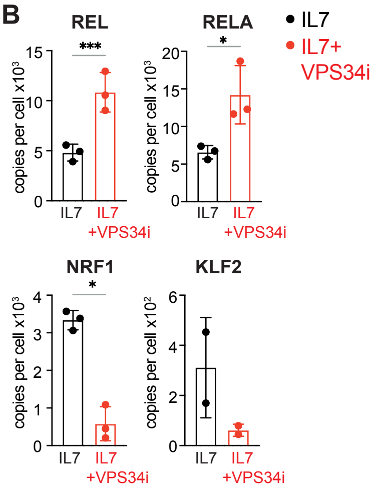

**Supplemental Figure S1**

**S1A,B** Quantitative proteomics data showing mean protein copy number per cell and RNASeq data shown as FPKM from naïve, 24 hour TCR activated P14 CD8 T cells and IL-2 maintained effector CD8 T cells (CTL) for **S1A)** GABARAP, GABARAPL2, ATG5, ATG7 and WIPI2 and **S1B)** FUNDC1, TAX1BP1, FAM134B, SEC63, ATL3 and TEX264.

**S1C** Representative flow cytometry data showing mCherry (left) and GFP (right) fluorescence profiles of ex vivo, naïve CD8 cells and IL-2 maintained effector CD8 T cells (CTL).

Proteomic data (S1A,B) are from Howden et al [1], all proteomic data are available on Immpres.co.uk [2]. FPKM mRNA data from Spinelli *et al.* [3] (S1A,B). All data are from a minimum of 3 biological replicates. Data points in bar charts are indicative of biological replicates. Error bars represent the mean +/- SD. P ≤ 0.0001 indicated by ****; P ≤ 0.0005 = ***; P ≤ 0.001 = **; P ≤ 0.05 = *

**Supplemental Figure S2**

**S2A** Quantitative proteomics data of the indicated proteins showing mean protein copy number per cell (right) and RNASeq data shown as FPKM (left) from CTL maintained with or without IL-2 over 24h.

**S2B** Quantitative proteomics data of the indicated proteins showing mean protein copy number per cell from CTL maintained with IL-2 , with IL-12+IL-18 , or no cytokine for 24h.

**S2C** Quantitative proteomics data of the indicated proteins showing mean protein copy number per cell (right) and RNASeq data shown as TPM (left) from CD8 T cells expanded with IL-2 (CTL) or expanded with IL-15 (memory-like).

The data in S2Aare from [3]; S2B are available on impress.co.uk [2]; S2C are from [4]. Data points are indicative of biological replicates.

**Supplemental Figure S3**

**S3A** Quantitative proteomics data showing mean protein copy number per cell of the indicated proteins from naïve or TCR activated CD8 24h.

**S3B** Quantitative proteomics data showing mean protein copy number per cell of the indicated proteins from CTL maintained with IL-2 , with IL-12+IL-18 , or no cytokine for 24h.

**S3C** Quantitative proteomics data showing mean protein copy number per cell of the indicated proteins from CD8 T cells expanded with IL-2 (CTL) or expanded with IL-15 (memory-like).

Proteomic data, S3A are from Howden et al [1], S3B are available on Immpres.co.uk [2], and S3C are from [4]. Data points in bar charts are indicative of biological replicates. Error bars represent the mean +/- SD.

**Supplemental Figure S4**

**S4A** Flow cytometry histograms (left) and MFI (right) of ribosomal protein S6 phosphorylation (pS6) in CTL maintained in amino acid replete media (RPMI) with or without rapamycin (rapa, 20 nM), in amino acid depleted media (HBSS) or in glucose free RPMI (no glucose) for 18hr.

**S4B** Flow cytometry histograms (left) and MFI (right) of ACC phosphorylation (pACC) in CTL maintained in regular, glucose containing media (RPMI) or in glucose free RPMI (no glucose) for 18hr.

Data points in bar charts are indicative of biological replicates.

**Supplemental Figure S5**

**S5A** Clustered enrichment analysis on proteins from amino acid replete (RPMI) CTL that were significantly increased with VPS34i treatment. The full enrichment table is available in Supplementary data file S5.

**S5B** Quantitative proteomics data showing mean protein copy numbers per cell of SQSTM1, GABARAPL2 and MAP1LC3B.

Data points in bar charts are indicative of biological replicates. Error bars represent the mean +/- SD. $P \leq 0.0001$ indicated by ****; $P \leq 0.0005$ = ***; $P \leq 0.001$ = **; $P \leq 0.05$ = *

**Supplemental Figure S6**

**S6A** Summed protein copy number of Complex I-V proteins of the mitochondrial electron transport chain (annotations: MitoCarta3.0 ‘OXPHOS’).

**S6B** Basal ECAR of CTL maintained in amino acid replete media (RPMI) or amino acid deprived (HBSS) with or without VPS34i for 18hr.

**S6C** Basal and oligomycin treated OCR measurements of CTL maintained in amino acid replete media (RPMI) or amino acid deprived (HBSS) with or without VPS34i for 18hr.

Data points in bar charts are indicative of biological replicates. Error bars represent the mean +/- SD. $P \leq 0.0001$ indicated by ****; $P \leq 0.0005$ = ***; $P \leq 0.001$ = **; $P \leq 0.05$ = *

**Supplemental Figure S7**

**S7A** MRPL20 and MRPL24 expression as a percentage of all large mitochondrial ribosome subunits.

**S7B-F** Protein copy numbers per cell of ERAL1 (S7B); proteins involved in mitochondrial transport (S7C); transferases (S7D); SLC7A3 (S7E) and transcriptional regulation (S7F).

Data points in bar charts are indicative of biological replicates. Error bars represent the mean +/- SD. P ≤ 0.0001 indicated by ****; P ≤ 0.0005 = ***; P ≤ 0.001 = **; P ≤ 0.05 = *

**Supplemental Figure S8**

**S8A** Mean protein copy numbers of BAX and BAK1 from naïve and OT1 CD8 T cells activated with SIINFEKL for the indicated times. Data available on Immpres.co.uk [2].

**S8B** Mean protein copy numbers of REL, RELA, NRF1 and KLF2 from CD8 T cells maintained in IL-7 with or without VPS34i.

Data points in bar charts or violin plots are indicative of biological replicates. Error bars represent the mean +/- SD. $P \leq 0.0001$ indicated by ****; $P \leq 0.0005$ = ***; $P \leq 0.001$ = **; $P \leq 0.05$ = *

**Supplemental Figure References**


1. Howden, A. J. M. *et al.* Quantitative analysis of T cell proteomes and environmental sensors during T cell differentiation. *Nature immunology* **20**, 1542–1554 (2019).

2. Brenes, A. J., Lamond, A. I. & Cantrell, D. A. The Immunological Proteome Resource. *Nat. Immunol.* **24**, 731–731 (2023).

3. Spinelli, L., Marchingo, J. M., Nomura, A., Damasio, M. P. & Cantrell, D. A. Phosphoinositide 3-Kinase p110 Delta Differentially Restrains and Directs Naïve Versus Effector CD8+ T Cell Transcriptional Programs. *Front Immunol* **12**, 691997 (2021).

4. Marchingo, J. M., Spinelli, L., Pathak, S. & Cantrell, D. A. PIM kinase control of CD8 T cell protein synthesis and cell trafficking. *bioRxiv* 2024.03.25.586560 (2024) doi:10.1101/2024.03.25.586560.
